# Sex differences in pediatric EEG inter-subject correlation during naturalistic movie watching: a large-scale characterization of the Healthy Brain Network EEG dataset

**DOI:** 10.64898/2026.09.07.749616

**Authors:** Eklavya Tomar, Aimar Silvan, Eugene Yujun Fu, Yong Xiong, Chi-wai Do

## Abstract

Inter-subject correlation (ISC) of EEG during naturalistic viewing is increasingly used as a candidate pediatric biomarker, yet its behavior at cohort scale remains uncharacterized. We characterize ISC in 1143 children and adolescents (765 male, 378 female, ages 5 to 21) from the Healthy Brain Network EEG dataset viewing four naturalistic film clips. Three results follow. First, ISC separates narrative from abstract stimuli by 1.7 to 2.4 fold and exceeds a resting-state baseline by 42 to 99 fold, and the previously reported developmental decline replicates throughout. Second, ISC is sex-dependent: males exceed females in every movie (*t* = 9.7 to 13.5; Cohen’s *d* = 0.61 to 0.85), concentrated in delta and theta, peaking at a frontocentral cluster and at ages 11 to 14. The direction matches an earlier report where the effect was marginal, and we resolve structure that sample could not. It survives three demographic and clinical robustness analyses, same-sex and size-matched templates, ocular component removal, and exclusion of the releases overlapping that report, and is attenuated, by approximately 10 percent adjusting for interpolation count and by 23 percent in recordings free of cluster interpolation, but not explained by differential channel interpolation. Evoked response magnitude, previously proposed as an alternative account, does differ by sex here (*d* = 0.55 to 0.60) but mediates only 8.6 to 18.1 percent of the effect. Third, ISC predicts none of four CBCL bifactor dimensions after FDR correction at any feature resolution: zero of 16 grand-mean, 2064 per-channel and 16 cluster tests, with confidence intervals excluding standardized associations beyond |*β*| = 0.106.

**Significance Statement:** Watching a film drives similar brain activity in different viewers, and how similar that activity is has been proposed as a marker of attention and of clinical status in children. We measured this in 1143 children and adolescents, roughly an order of magnitude more than previous work. Boys showed more similar responses than girls, an effect largest in slow frontocentral activity and peaking around ages 11 to 14. This confirms a previously tentative finding and excludes or bounds several measurement explanations for it, including the one the original authors raised but could not settle, which we find accounts for under a fifth of the difference. Against this, the same measure carried no detectable information about parent-reported psychopathology, and these data exclude all but the smallest of the associations such studies typically report. The measure is developmentally and sex-sensitive but, on this protocol, clinically uninformative.

## 1 Introduction

Naturalistic stimuli such as films and narratives engage the brain in ways more representative of everyday cognition than the brief, repetitive stimuli of classical paradigms, recruiting attention, memory, language, social cognition, and emotion in ecologically valid configurations [1]–[3]. The dominant analytic approach to such data is inter-subject correlation (ISC), the cross-subject correlation of neural time series during shared stimulus viewing [1], [5]. ISC isolates the shared, stimulus-driven component of the response and tracks attention, engagement, narrative comprehension, and memory [4], [6]. Because it requires no model of stimulus features, only that subjects view the same content at the same time, it can be computed on any sufficiently long shared-stimulus recording.

Naturalistic stimuli are particularly attractive in pediatric work, since children sustain attention to engaging films in ways they cannot to abstract laboratory tasks [7], [8]. The foundational study of pediatric EEG ISC, by Petroni and colleagues [9], reported two findings across two cohorts. Neural response variability increased with age, consistent with developmental refinement toward greater individuation. And among children, male participants showed less variable responses than female participants, equivalently higher ISC, which the authors read as indicating that male brains are less neurally mature in this window. The sex effect was marginal in both cohorts once age was partitioned (*p* = 0.05 at *n* = 114; *p* = 0.1 at *n* = 303), and the authors noted that it did not survive control for the magnitude of the visual evoked response, leaving open whether it reflects shared processing or simply stronger stimulus-evoked signal in males.

ISC has also attracted attention as a candidate clinical biomarker. Reduced ISC has been shown in adults with attention-deficit/hyperactivity disorder using fMRI [12], and EEG functional connectivity from naturalistic viewing carries phenotypic signal at scale [13]. The natural extension asks whether pediatric ISC predicts dimensional psychopathology of the kind measured by the Child Behavior Checklist [14] and its bifactor decomposition [15], [16]. Recent benchmarking sharpens the question: the 2025 NeurIPS EEG Foundation Challenge [17] set over 1100 teams and participants to predict CBCL externalizing scores from HBN-EEG recordings, and only three finished below the organizers’ threshold for improvement on a mean-prediction baseline [18].

Two developments make a decisive test feasible. The HBN-EEG release [19], derived from the Healthy Brain Network biobank [20], provides high-density recordings from a large pediatric cohort during four naturalistic clips. And automated bad-channel detection [21] and machine-learning component classification [22] make principled large-cohort preprocessing tractable in ways it was not when the foundational work was done.

No prior study has reported developmental or sex effects on pediatric EEG ISC at samples exceeding 1000, tested its relationship to dimensional psychopathology at that scale, or resolved the spectral, topographic and developmental structure of the sex effect. Using 1143 subjects from the first eight HBN-EEG releases, we characterize ISC magnitude and topography, replicate the developmental decline, and confirm and resolve the sex effect. We evaluate its robustness through three demographic and clinical analyses, a check on cohort overlap with the original report, three analyses targeting the ISC computation and the measurement itself, and a direct reproduction of the evoked-magnitude control that the original report raised but could not settle. Finally we test whether ISC predicts any of four CBCL bifactor dimensions at three levels of feature resolution.

This paper makes three contributions. The first is characterization: the magnitude, topography, stimulus dependence and developmental trajectory of pediatric EEG ISC at a sample size an order of magnitude beyond prior work, with a resting-state negative control separating stimulus-driven from spurious shared variance. The second is confirmation and resolution of the sex effect: recovered at *n* = 1143, against original cohorts of 114 and 303, and under a different ISC estimator, resolved into its delta concentration, frontocentral weighting and inverted-U developmental trajectory, with the previously proposed evoked-magnitude mechanism quantified at 8.6 to 18.1 percent of the total. The third is a bound: ISC predicts none of four CBCL bifactor dimensions at controlled false discovery rate across three feature resolutions, with confidence intervals excluding standardized associations larger than |*β*| = 0.106.

## 2 Methods

### 2.1 Participants

We analyzed data from the Healthy Brain Network EEG (HBN-EEG) release [19], a Findable, Accessible, Interoperable, Reusable (FAIR) public release of high-density EEG data from a community-referred pediatric cohort recruited by the Child Mind Institute. The full HBN study enrolls children and adolescents aged 5 to 21 years from the New York metropolitan area with a parent-reported concern about a child’s mental health, learning, or behavior; recruitment is community-referred rather than clinic-based, and because families are invited on the basis of perceived clinical concern, the sample is expected to contain a high proportion of participants affected by psychiatric illness [20]. The study design does not include a separately recruited comparison group. All data analyzed here derive from the public HBNEEG release. Informed consent and assent were obtained from participants and their legal guardians by the Child Mind Institute under its own institutional review board approval, as described in the source dataset publications [19], [20]. The present work is a secondary analysis of deidentified, publicly released data involving no new data collection and no participant contact, and therefore required no additional ethical approval.

Sex is recorded in the public HBN-EEG release as a binary field in `participants.tsv`, reported at enrollment by a parent or guardian. It is not accompanied by a separate gender identity field, by information on pubertal stage, or by any hormonal measure. We therefore use the term sex throughout, refer to the two recorded categories as male and female, and note that this field cannot distinguish sex assigned at birth from gender identity in participants for whom these differ.

The HBN-EEG release we analyzed comprises eight data releases (R1 through R8), made available in BIDS format on the FCP-INDI Amazon S3 bucket. For each release, we identified the subset of participants with complete Child Behavior Checklist (CBCL) bifactor scores in `participants.tsv` across all four dimensions of interest (general psychopathology factor, attention, internalizing, externalizing). Among these participants, we further restricted to subjects who had usable EEG recordings for all four naturalistic movie-watching tasks. Subjects with truncated stimulus presentation (defined post hoc as recording duration shorter than 200 seconds for The Present, due to a stimulus delivery issue identified in a small subset of R5 and R8 subjects) were excluded from the corresponding movie’s analyses.

Final analytic cohort size after preprocessing and quality control was *n* = 1143 subjects (765 male, 378 female; sex ratio 2.02:1) with mean age 10.18 years (range 5.0 to 21.8). Cohort breakdown by release: R1 (*n* = 117), R2 (*n* = 114), R3 (*n* = 138), R4 (*n* = 259), R5 (*n* = 207), R6 (*n* = 48), R7 (*n* = 124), R8 (*n* = 136). The male skew is largely a property of the source cohort: the HBN-EEG release is 35 percent female across its first nine releases [19], against 33.1 percent in the present analytic cohort drawn from the first eight, so quality control and the requirement of usable recordings for all four movies shifted the ratio slightly further towards males. Alexander et al. [20] attribute the skew in the parent cohort principally to the prevalence of ADHD, commonly estimated at a 3:1 male to female ratio in children. Demographic and psychopathology characteristics of the cohort are summarized in Table 1.

**TABLE 1.** Cohort composition by HBN release. The analytic cohort comprises the subset of subjects passing loose QC on all four movies with complete CBCL bifactor scores.

| Release | Available | QC pass | Pass rate | Cumulative |
| --- | --- | --- | --- | --- |
| R1 | 120 | 117 | 97.5% | 117 |
| R2 | 120 | 114 | 95.0% | 231 |
| R3 | 157 | 138 | 87.9% | 369 |
| R4 | 293 | 259 | 88.4% | 628 |
| R5 | 282 | 207 | 73.4% | 835 |
| R6 | 102 | 48 | 47.1% | 883 |
| R7 | 247 | 124 | 50.2% | 1007 |
| R8 | 214 | 136 | 63.6% | 1143 |
| Total | 1535 | 1143 | 74.5% | 1143 |

Cohort summary: Sex: 765 M / 378 F (ratio 2.02:1) Age: mean 10.18 y, range 5.0 to 21.8

### 2.2 Stimuli

Each subject viewed four short film clips during a single recording session, in counterbalanced order across subjects. The clips were selected by the Child Mind Institute to span a range of narrative and visual content types relevant to pediatric attention and engagement studies:

- *Despicable Me* (170.5 s): an excerpt from the animated feature film, depicting a comedic interaction with three children and animated kittens. Animated, narrative content with dialog.
- *Diary of a Wimpy Kid* (117.4 s): the theatrical trailer for the 2010 live-action adaptation, featuring rapid scene changes and live-action footage with cartoon overlays.
- *Fun with Fractals* (163.0 s): an educational mathematical animation depicting iterative fractal patterns set to instrumental music, with no narrative or speech content.
- *The Present* (203.1 s): an Academy Award-shortlisted animated short film by Jacob Frey (2014), depicting an emotionally arc-driven story between a boy and a three-legged puppy. Animated, narrative content with minimal dialog.

Three of these clips (Despicable Me, Diary of a Wimpy Kid, Fun with Fractals) were also used by Petroni et al. [9], at matched duration for the first two. Their version of Fun with Fractals ran approximately 274 s against 163.0 s here, so absolute ISC magnitudes for that stimulus are not directly comparable between the two studies.

Source video files for these stimuli were reconstructed from original distribution materials by collaborators on the HBN-EEG release to address timing irregularities in the stimulus delivery system that had been documented through prior eye-tracking validation. Reconstructed videos were upsampled to 30 fps and frame-aligned to recorded EEG via the stimulus onset and offset markers in the BIDS `events.tsv` files. The nominal playback duration for The Present is 203.133 seconds under the original presentation specification; recordings shorter than 200 seconds were excluded and the remainder cropped to their common minimum duration as described in Section 2.6.

### 2.3 EEG Data Acquisition

EEG data were acquired by the HBN study team in a sound-shielded room using a 128-channel EGI HydroCel Geodesic Sensor Net at a sampling rate of 500 Hz, with an acquisition bandpass of 0.1 to 100 Hz and the Cz electrode as the recording reference [20]. Data runs not recorded at 500 Hz were resampled to that rate during preparation of the public release [19]. The recorded montage comprises 128 scalp electrodes plus the Cz reference; following common average re-referencing (Section 2.4) the reference channel is recovered, giving the 129 channels used throughout the analyses. We analyzed data as released, without modification to the BIDS-formatted source files.

### 2.4 Preprocessing

We developed and validated a preprocessing pipeline (designated `v3concat`) tailored to the short-clip naturalistic stimulus structure of HBN movie tasks. The pipeline was implemented in Python 3.11 using MNE-Python (version 1.12) [23], PyPREP [21], and MNE-ICALabel [22].

For each subject, all four movie recordings were processed jointly in a single pipeline instance to maximize the data available for independent component analysis (ICA) decomposition. Per-subject processing proceeded as follows. **Stimulus-window cropping**. Each movie recording was cropped to the [video start, video stop] window defined by stimulus markers in `events.tsv`, removing pre-task baseline, inter-trial intervals, and post-task data.

#### Line noise removal

Notch filters were applied at 60 Hz and 120 Hz using MNE-Python defaults to suppress power line and harmonic contamination characteristic of US-recorded data.

#### Dual-stream filtering

Two parallel filtered copies of each recording were generated. The first (`raw_ica`) was bandpass filtered 1 to 100 Hz and used for ICA fitting. The second (`raw_analysis`) was bandpass filtered 1 to 20 Hz and used for downstream ISC analyses. The 1 Hz high-pass minimizes ICA convergence issues from slow drift; the 20 Hz low-pass on the analysis stream excludes high-frequency muscle artifact while retaining the frequency range of interest for movie-evoked synchronization.

#### Common average reference

Both filtered streams were re-referenced to the common average across all recorded channels, which recovers the Cz reference electrode and gives the 129 channels used throughout. This step preceded bad-channel detection, so the average includes channels subsequently flagged as bad. PyPREP’s robust referencing procedure, which iteratively excludes bad channels from the reference before detection, was not used; the consequences are considered in Section 4.5.

#### Bad channel detection

PyPREP’s `NoisyChannels` routine [21] was then applied separately to each movie’s average-referenced `raw_ica` copy, flagging channels by criteria including deviation from the median, low correlation with neighbors, low signal-to-noise ratio, and RANSAC-based prediction failure. Flagged channels were spherically interpolated in both streams using MNE-Python’s standard interpolation routine. Because detection follows referencing, the recovered Cz channel is screened on the same footing as a recorded electrode.

#### Independent Component Analysis

The four cropped, filtered `raw_ica` streams for a given subject were concatenated using MNE’s `concatenate_raws` function, which inserts a boundary annotation at each junction so that filtering and decomposition do not span clips. ICA was then fit on the concatenated stream using the infomax extended algorithm [24] with 20 components and a maximum of 500 iterations. The decomposition was restricted to 20 components rather than the full channel rank, a substantial dimensionality reduction on 129 channels. We did not optimize this value and make no claim that 20 is preferable to another choice. Its consequences are measured rather than assumed: the reproducibility of the resulting decomposition across random initializations is quantified in Section 2.10, and the robustness of the sex effect to re-fitting the decomposition per subject is established in Section 3.4.3. Concatenation across the four movies provided approximately 10 minutes of EEG data per subject for ICA fitting, substantially more than would be available from any single movie of 2 to 3 minutes alone. Per-clip ICA, by contrast, would fit each decomposition on 2 to 3 minutes of data, approximately one quarter of the data available to the concatenated fit, so the concatenated fit was a deliberate design choice rather than a neutral alternative; boundary annotations prevented the decomposition from modeling cross-clip discontinuities.

#### Component classification

The fitted ICA decomposition was passed through MNE-ICALabel [22], which classifies each component into one of seven categories (brain, muscle, eye blink, heart, line noise, channel noise, other) using a deep learning model trained on adult and pediatric EEG. Components classified as muscle, eye blink, heart, line noise, or channel noise were marked for removal regardless of classifier confidence. Median classifier confidence across all 20 components was retained as a quality control metric. Because the decomposition is fitted once per subject, this is a per-subject value recorded identically for each of that subject’s four recordings. The same exclusion list was applied to all four movies for that subject.

#### Signal reconstruction

The marked components were removed by applying `ICA.apply()` separately to each movie’s `raw_analysis` stream. Following IC removal, each movie was re-referenced to the common average a second time and resampled to 200 Hz to standardize temporal resolution across analyses.

The `v3concat` pipeline was developed iteratively, with two earlier iterations (v1, v2) tested on a subset of subjects before convergence. A single-subject pilot comparing `v3concat` output to per-movie ICA fits on the same data is reported in Supplementary Materials; agreement was broad at the typical channel but not uniform across channels, and the joint fit additionally provides a single uniform exclusion list per subject.

### 2.5 Quality Control

Quality control was applied after preprocessing. Three metrics were computed: (i) median per-channel standard deviation in microvolts (`std_p50`), a measure of overall signal magnitude after artifact removal; (ii) median ICALabel classification confidence across all 20 components (`icalabel_conf`); and (iii) the number of components classified as brain (`ic_brain`), a measure of usable neural signal capacity in the decomposition. Only (i) is computed per recording; because the decomposition is fitted once per subject on the concatenated stream, (ii) and (iii) are per-subject values that take the same value for each of that subject’s four recordings.

Two QC tiers were defined. Loose QC, used for the primary analyses: `std_p50` *≤* 40 *µ*V, `icalabel_conf` *≥* 0.50, `ic_brain` *≥* 3. Strict QC, used for sensitivity analyses: `std_p50` *≤* 20 *µ*V, `icalabel_conf` *≥* 0.65, `ic_brain` *≥* 6. Subjects had to pass loose QC on all four movies for inclusion in the primary cohort. Pass rates by release ranged from 47.1% (R6) to 97.5% (R1), with notably lower rates in R6 and R7 reflecting documented variability in HBN data quality across releases. Statistical analyses controlled for release as described in Section 2.7.

### 2.6 Inter-Subject Correlation Computation

For each of the four movies and each subject in the cohort, leave-one-out (LOO) inter-subject correlation was computed following the standard procedure [4], [9]. For each channel and each subject, the EEG time series was correlated (Pearson) with the average time series across all other subjects in the cohort for that channel and that movie. The result is a per-subject, per-channel ISC value. Subject-level grand-mean ISC was computed as the mean across the 129 channels.

This estimator differs from that of Petroni et al. [9], who computed ISC on the projections of the three maximally correlated components recovered by correlated component analysis [5]. Their measure is spatially filtered and dimensionality-reduced; ours is computed in channel space and averaged. Absolute magnitudes are therefore not comparable between the two studies, and only the direction and relative ordering of effects should be compared.

Time series alignment was achieved by cropping all subjects to the common minimum recording duration for each movie. Discrepancies between subjects in recording length, where present, were minor (within 0.5 seconds for 99.9% of recordings). For The Present, the nominal presentation duration of 203.133 seconds served as the reference against which truncated recordings were identified and excluded (Section 2.1); retained recordings were then cropped to their common minimum duration in the same way as the other three movies.

ISC was computed on the broadband (1 to 20 Hz) signal for the primary cohort analyses. For the frequency-band decomposition (Section 2.8), the same procedure was applied to band-limited signals.

### 2.7 Statistical Analyses

#### Primary regression

The primary statistical model was the per-movie ordinary least squares regression:

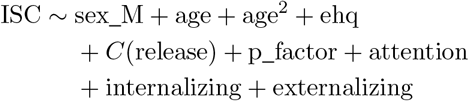

where `sex_M` is a binary indicator for male sex, `age` is in years, `ehq` is the Edinburgh Handedness Questionnaire total score, *C*(release) is a categorical fixed effect for HBN release (R1 through R8, allowing release-specific intercepts), and the four CBCL factors are entered as continuous covariates. The `sex_M` coefficient is the primary effect of interest; the regression was fit separately for each of the four movies. Eleven subjects (10 male, 1 female) missing one or more model covariates were removed by listwise deletion, giving per-movie regression samples of 1132, and 1128 for The Present. Because age is entered untransformed alongside age^2^, the linear coefficient in this model is the fitted slope at age zero, five years below the youngest participant, and is not interpretable as the effect of age within the sample. The developmental effect reported in Section 3.3 is therefore taken from the same model with the quadratic term removed, while the sex coefficient is reported from the model as specified above, in which age^2^ serves as a flexible control. Statistical significance was assessed at *α* = 0.05. The prespecified criterion for treating the sex effect as robust to demographic confounding was significance at *p <* 0.05 in at least 3 of the 4 movies in the fully adjusted regression. The observed result exceeded that criterion, reaching *p <* 0.001 in all four (Table 4).

#### Multiple comparisons

Where multiple tests were conducted across movies or CBCL factors, Benjamini-Hochberg false discovery rate (FDR) correction [26] was applied at *q <* 0.05. Correction families were defined per analysis: across the four movies for per-movie effects, across the 129 channels within each movie for the per-channel sex effect, and jointly across all 2064 tests for the per-channel CBCL analysis. Correcting jointly across all 2064 tests is the more stringent of these options. Because that analysis reports a null, the stringency works against the strength of the claim rather than for it: a less stringent family would make individual hits easier to obtain, and their absence correspondingly more informative. We therefore do not rest the null on the correction. The bound in Table 9 is computed from confidence intervals and does not depend on the choice of family, and the sensitivity of the per-channel null to the partition was additionally checked post-hoc under two less stringent families, reported with the per-channel results.

#### ISC × CBCL

Each of four CBCL factors (general psychopathology, attention, internalizing, externalizing) was tested against grand-mean ISC for each of four movies, yielding 16 tests. Each test fitted a regression of the form ISC ~ cbcl factor + sex M + age + age^2^ + ehq + *C*(release), with the `cbcl_factor` coefficient as the effect of interest. FDR correction was applied across all 16 tests. Because this analysis reports a null, we additionally report each coefficient in standardized units with its 95% confidence interval, and the minimum detectable effect at 80% power for the achieved standard error, so that the result functions as a bound rather than an absence of evidence.

#### Per-channel sex effect

For the topographic analyses, per-channel ISC values were tested for sex differences using independent samples t-tests at each of 129 channels, with FDR correction across channels within each movie.

#### Interaction tests

The sex × movie interaction was tested using a single regression on the long-format data (one row per subject-movie observation) with `sex_M`, *C*(movie), and their interaction, plus age, ehq, and *C*(release) as covariates; significance was assessed via an F-test on the joint set of interaction terms compared to the no-interaction reduced model. The sex *×* age interaction was tested per-movie using `sex_M` *×* `age_centered` (linear) and `sex_M` *×* `age_centered`^2^ (quadratic) terms.

#### Robustness analyses

The sex effect was further evaluated through two prespecified robustness analyses: (1) age stratification (5 age bins, sex effect tested per bin per movie); and (2) strict QC subset analysis (*n* = 486). Both are reported in the Results regardless of outcome. Two further analyses were added post-hoc: a low-symptom subset described in Section 3.4.2, addressing potential clinical-sample confounding, and a release-exclusion sensitivity analysis addressing partial cohort overlap with the replication sample of Petroni et al. [9]. A post-hoc analysis on differential channel interpolation between sexes is described in Section 3.4.3. **Resting-state baseline**. As a methodological control, ISC was computed using identical procedures on R1 RestingState recordings (*n* = 132, recording durations 5.7 to 16.8 minutes per subject; cropped to a common length for ISC computation). This served as a negative control to verify that the LOO procedure does not produce spurious ISC in the absence of a shared stimulus.

### 2.8 Frequency-Band Decomposition

To localize the sex effect to a specific neural frequency band, ISC was recomputed in four bands of interest after additional bandpass filtering of the preprocessed signals: delta (1 to 4 Hz), theta (4 to 8 Hz), alpha (8 to 12 Hz), and beta (12 to 20 Hz). Filter design was an FIR bandpass filter (firwin design, zero-phase) applied via MNE-Python’s `filter_data` function with default parameters. For the delta band this yielded a Hamming-windowed filter of 661 samples with lower and upper transition bandwidths of 1.00 and 2.00 Hz, placing the −6 dB cutoffs at 0.50 and 5.00 Hz. Because the analysis stream had already been high-pass filtered at 1 Hz with an identical 1.00 Hz transition, the two roll-offs coincide, so content between 0.5 and 1 Hz is attenuated twice, by approximately 12 dB at 0.5 Hz. The delta passband above 1 Hz is unaffected. Delta results should therefore be read as covering 1 to 4 Hz with a cascaded roll-off below 1 Hz.

Because per-band computation is more memory-intensive than broadband ISC, this analysis was performed on a balanced subsample of *n* = 400 (200 male, 200 female; random sample within sex with seed 42) drawn from the full QC-pass cohort, of which 398 have usable recordings for The Present. The full preprocessing chain was applied identically; only the additional bandpass filter was different across band conditions. Leave-one-out templates for the band-limited analyses were formed within this subsample rather than from the full cohort, so band ISC magnitudes are not directly comparable to the full-cohort broadband values. Sex effects within each band were tested using the same primary regression structure as Section 2.7.

### 2.9 Evoked Response Magnitude

Petroni et al. [9] raised the possibility that the sex difference in ISC reflects a difference in the magnitude of the stimulus-evoked response rather than in the extent to which responses are shared across subjects. Because ISC is a correlation, a subject whose evoked response is large relative to background neural activity will show higher ISC without any difference in how stereotyped that response is. They tested this using steady-state visual evoked potential (SSVEP) magnitude from a flickering-grating stimulus, and reported that the sex effect on ISC did not survive the control, while cautioning that the SSVEP subsample was smaller than the full cohort (*N* = 84 against 114).

The HBN protocol includes a Surround Suppression task using the same paradigm [11] at the same 25 Hz flicker frequency, which allows this control to be reproduced directly. Foreground disks flicker on and off at 25 Hz against a static surround, presented in 2.4 s trials at four foreground contrast levels (0%, 30%, 60%, 100%) crossed with surround conditions, across two runs per subject.

We downloaded Surround Suppression recordings for the balanced *n* = 400 subsample used for the band decomposition; 354 of the 400 have these recordings available in the public release. Processing was deliberately minimal, since SSVEP magnitude is a narrowband amplitude measurement rather than a source-separation problem: notch filtering at 60 and 120 Hz, bandpass filtering 1 to 45 Hz (the 1 to 20 Hz analysis stream used elsewhere excludes 25 Hz entirely and cannot be used here), and common average re-referencing. No ICA was applied.

Trials were epoched from stimulus onset, with the first 200 ms discarded to remove the onset visual evoked potential, giving a 2.2 s analysis window. At this window length 25 Hz falls exactly on an integer discrete Fourier transform bin, so no taper was applied. Trials were rejected using a robust global criterion: the median across channels of per-channel root mean square amplitude, thresholded at 3 median-absolute-deviation-scaled standard deviations in log space, with an absolute median peak-to-peak ceiling of 250 *µ*V as a backstop and a hard cap of 25 percent of trials per recording. Criteria based on whether any single channel exceeds a threshold were avoided: with 129 channels and no prior artifact removal, such rules are triggered by any persistently bad electrode on essentially every trial, and bad-channel counts differ by sex in this cohort (Section 3.4.3), so that failure mode would bias the covariate on the variable under test. Median trial retention was 87.5 percent, and no subject was excluded by these criteria: all 354 subjects with available recordings yielded usable data.

For each subject we computed the amplitude at 25 Hz divided by the mean amplitude in neighboring bins, giving a signal-to-noise ratio at the flicker frequency. Analysis electrodes were the five with the highest group-mean SNR, selected on the group average and applied uniformly to every subject. Three measures were derived: an incoherent average, matching the per-trial averaging used by Petroni et al.; a coherent average, in which trials are averaged in the time domain before transformation, which exploits the phase-locking of the SSVEP and yields a more reliable per-subject estimate; and a coherent average restricted to foreground contrast at or above 60%. The incoherent measure is used as primary for comparability with the original report, and the other two are reported as robustness.

The sex effect on ISC was then re-estimated with SSVEP SNR added to the primary regression (Model 1) and with SSVEP SNR plus quality-control covariates added (Model 2). We additionally report a formal mediation decomposition of the sex effect into a path running through SSVEP magnitude and a remaining direct path, with bootstrap confidence intervals on the indirect path (2000 resamples, seed 42).

### 2.10 Decomposition Stability

To quantify what the concatenated fit contributes, we compared per-clip and concatenated ICA using the cluster quality index *I*_*q*_ [25] on 30 subjects with 20 random initializations per condition, together with two duration-matched controls separating sample volume from content heterogeneity. Concatenation improves component reproducibility by a reliable margin (mean *I*_*q*_ 0.759 against 0.701 across the four per-clip fits, paired *t* = 8.98, *p* = 7.2 *×* 10^−10^), raising components meeting the conventional stability threshold from roughly 5 of 20 to 8.6 of 20. It does not resolve non-convergence, and clip duration does not predict stability across the 117 to 200 s range used here, so the gain reflects content heterogeneity rather than clip duration as such; volume is not irrelevant, since the full concatenation exceeds its duration-matched control. The procedure and full per-condition results are reported in Supplementary Materials.

### 2.11 Software and Reproducibility

All analyses were performed in Python 3.11. Primary dependencies were MNE-Python (1.12), PyPREP (0.6), MNE-ICALabel (0.8), pandas (3.0), numpy (2.4), scipy (1.17), scikit-learn (1.8), and statsmodels (0.14). Figures were generated using matplotlib (3.10). All preprocessing and analysis code needed to reproduce the reported results, together with an environment specification and documentation of the run order from raw data to the tables and figures, is publicly available at https://github.com/EklavyaT/hbn-isc. Disclosure of AI assistance in the preparation of this work is given in the Acknowledgment. The HBN-EEG dataset itself is publicly available without registration via the FCP-INDI S3 bucket as cited above and is not redistributed in the repository; the reported subsamples regenerate exactly from the deposited code under the fixed random seed.

Random seeds were fixed at 42 for every subsampling and resampling procedure in the paper. These are the band-decomposition subsample, the balanced subsample for the ocular re-fit, the size-matched template draws, the bootstrap confidence intervals on ISC magnitudes and on Cohen’s *d*, the strict-QC matched-draw control, the mediation bootstrap, and the removal controls and stratum bootstrap of Section 3.4.3. Two steps of the preprocessing pipeline are stochastic: the ICA initialization and the RANSAC stage of bad-channel detection. Both use a single fixed seed of 42 applied to every subject rather than a per-subject seed, so the pipeline is deterministic given the deposited code.

## 3 Results

### 3.1 Cohort Characteristics and ISC Magnitudes

The final analytic cohort comprised 1143 subjects passing loose QC on all four movies (765 male, 378 female; mean age 10.18 years, range 5.0 to 21.8; sex ratio 2.02:1). Cohort demographics broken down by release are presented in Table 1.

Grand-mean ISC values were comparable to those reported in prior pediatric naturalistic-stimulus EEG studies. Across the cohort, mean ISC ranged from 0.056 (Fun with Fractals) to 0.132 (Diary of a Wimpy Kid), with the three narrative-containing stimuli (Despicable Me, Diary of a Wimpy Kid, The Present) showing 1.7- to 2.4-fold higher ISC than the abstract mathematical animation (Fun with Fractals). Per-movie statistics are summarized in Table 2 and Figure 1.

**TABLE 2.** Inter-subject correlation magnitudes per movie. Mean grand-mean ISC across the cohort with 95% bootstrap confidence intervals (1000 resamples). Resting-state ISC computed on R1 RestingState recordings as a methodological baseline.

| Stimulus | $n$ | Mean ISC (95% CI) | SD |
| --- | --- | --- | --- |
| Despicable Me | 1143 | 0.097 (0.094, 0.100) | 0.046 |
| Diary of a Wimpy Kid | 1143 | 0.132 (0.128, 0.136) | 0.063 |
| Fun with Fractals | 1143 | 0.056 (0.054, 0.058) | 0.031 |
| The Present | 1139 | 0.112 (0.109, 0.115) | 0.051 |
| Resting state | 132 | 0.00133 (−0.001, 0.003) | 0.010 |

**Fig. 1.**
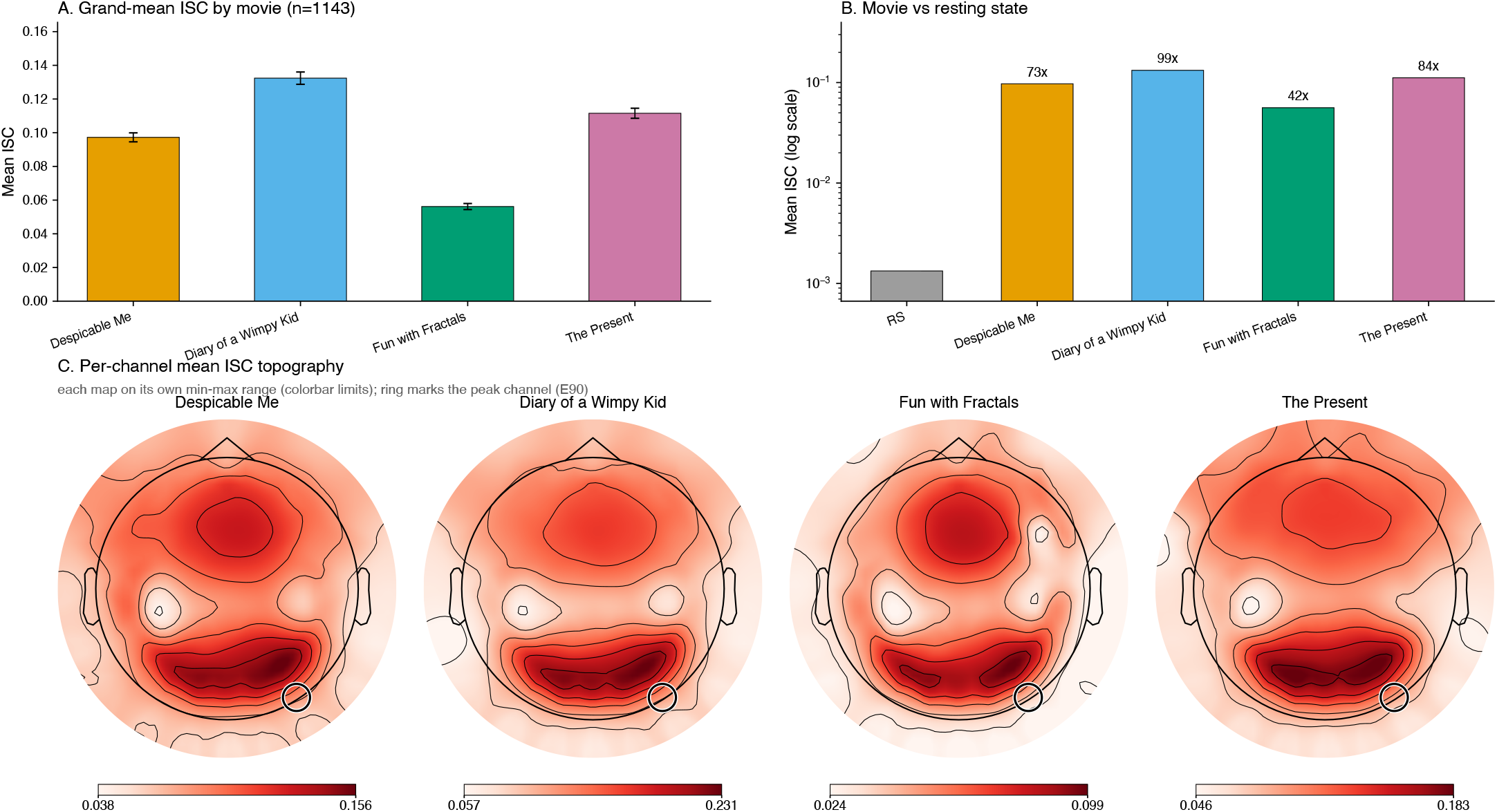
ISC magnitudes and topographic distribution. (A) Grand-mean ISC across all subjects for each of the four movies, mean *±*95% CI. The three narrative stimuli show 1.7 to 2.4*×* higher ISC than the abstract control. (B) Resting-state ISC (*n* = 132) compared to movie ISC on log scale, showing 42 to 99*×* magnitude separation that confirms the metric is stimulus-driven. (C) Channel-wise mean ISC for each movie, scaled per movie to that movie’s own range, which is printed beneath each map. ISC is high across the whole scalp with a parieto-occipital maximum at E90 in every movie, ringed; the posterior half of the montage exceeds the anterior half by 3 to 7 percent.

The narrative-vs-abstract distinction is consistent with the established pattern that narrative content drives stronger cross-subject neural synchronization than abstract or non-narrative stimuli [2], [4]. Differences across the three narrative movies (1.4-fold range) likely reflect differences in visual dynamics rather than narrative structure per se, consistent with prior reports that visual motion is the strongest single driver of EEG ISC in video viewing [27]. Channel-wise, ISC was high across the whole scalp and maximal parieto-occipitally in every movie, with E90 the highest-ISC channel throughout (Figure 1C). The posterior half of the montage exceeded the anterior half by only 3 to 7 percent in mean ISC, so the topography is a broad field with a posterior maximum rather than a localized cluster. Nine channels (E65, E70, E71, E75, E76, E83, E84, E90, E91) were in the top ten in all four movies, all of them posterior, and no anterior channel entered the top ten in any movie. The location of the maximum is consistent with the dominance of visual-cortical processing in determining where movie-evoked synchronization is largest.

Across the three stimuli shared with Petroni et al. [9], the rank ordering of ISC magnitude is the same in both studies (Diary of a Wimpy Kid *>* Despicable Me *>* Fun with Fractals), despite the different estimator, cohort, and preprocessing. Absolute values are not comparable for the reasons given in Section 2.6.

### 3.2 Resting-State Baseline

To verify that the ISC pipeline does not produce spurious shared variance in the absence of a stimulus, ISC was computed on R1 RestingState recordings. The resting-state cohort comprised *n* = 132 R1 subjects with usable resting-state recordings and complete CBCL scores; this differs from the R1 four-movie analytic cohort of *n* = 117 in Table 1 because resting-state inclusion did not require all four movies to be usable. Resting-state ISC was statistically indistinguishable from zero (mean = 0.00133, SD = 0.010, t-test against zero: *t* = 1.51, *p* = 0.13). Per-channel mean ISC ranged from − 0.011 to 0.006 across all 129 channels, with no channel exceeding 0.01. Movie ISC was 42-to 99-fold larger than resting-state ISC across the four movies, with the per-channel distributions of resting-state and movie ISC values entirely non-overlapping (resting-state per-channel maximum: 0.006; movie per-channel minimum: 0.024 in Fun with Fractals). Two-sample t-tests confirmed massive separation between the two conditions for every movie (*t* = 20.2 to 24.6, all *p <* 1 *×* 10^−78^). This validates that the LOO ISC procedure recovers shared variance only when subjects view a synchronized stimulus.

The resting-state baseline reported by Petroni et al. [9] for a different cohort under a different estimator was 0.001, effectively the same floor. The two studies therefore agree on the value of the negative control even though their movie ISC magnitudes are on different scales.

### 3.3 Developmental Effect on ISC

The age effect on ISC reported by Petroni et al. [9] replicated cleanly at substantially larger sample size. Age was a significant negative predictor of ISC in 4 of 4 movies (Table 3, Figure 2), all four at *q*_FDR_ < 0.001, with every confidence interval excluding zero. The effect held for Fun with Fractals as well as for the three narrative stimuli, indicating the developmental pattern is not specific to narrative content. Raw mean ISC fell by 33 to 41 percent from the youngest age bin (5 to 8) to the oldest (17 to 22) in every movie, so the decline is visible without adjustment. The oldest bin is sparse (20 male and 12 female subjects) and is excluded from the sex-difference analyses of Section 3.4.2 for that reason; it is retained here because a group mean over 32 subjects is a far more stable quantity than a between-sex contrast within them.

**TABLE 3.** Developmental effect on grand-mean ISC per movie, from the primary regression with the quadratic age term removed so that the coefficient is interpretable as the average linear slope across the sampled age range. Units are ISC per year. FDR correction is across the four movies. Direction is uniformly negative and every confidence interval excludes zero. The concavity of the trajectory is reported in the text.

| Movie | $\beta_{\text{age}}$ (95% CI) | $q_{\text{FDR}}$ |
| --- | --- | --- |
| Despicable Me | -0.00231 (-0.00313, -0.00149) | $1.5 \times 10^{-7}$ |
| Diary of a Wimpy Kid | -0.00246 (-0.00356, -0.00136) | $1.8 \times 10^{-5}$ |
| Fun with Fractals | -0.00125 (-0.00181, -0.00069) | $1.8 \times 10^{-5}$ |
| The Present | -0.00181 (-0.00269, -0.00094) | $5.2 \times 10^{-5}$ |

**TABLE 4.** Sex effect on grand-mean ISC per movie. Columns 2 to 4: unadjusted two-sample t-test (males vs females) with Cohen’s *d* (bootstrap 95% CIs computed with 1000 resamples all excluded zero; ranges: 0.541 to 0.807 for Despicable Me, 0.703 to 0.968 for Diary of a Wimpy Kid, 0.483 to 0.739 for Fun with Fractals, 0.715 to 0.982 for The Present). Columns 5 to 7: full regression with all controls (sex, age, age^2^, EHQ, release, four CBCL scores). Effect direction is uniformly positive (males *>* females) across all four stimuli at both unadjusted and fully-adjusted levels. Welch’s t-tests yield essentially identical results (*t* = +9.6 to +13.1; all Welch *p <* 1.3 *×* 10^−20^); Levene’s tests indicate variances are statistically equivalent in 3 of 4 movies (only The Present nominally unequal at *p* = 0.018, not surviving Bonferroni correction across 4 movies).

| Movie | Unadjusted |  |  | Adjusted |  |  |
| --- | --- | --- | --- | --- | --- | --- |
| | $t$ | $p$ | $d$ | $\beta$ | $p$ | $\eta_p^2$ |
| Despicable Me | 10.6 | 2.6e-25 | 0.67 | 0.0288 | 1.2e-24 | 0.090 |
| Diary of a Wimpy Kid | 13.2 | 3.1e-37 | 0.83 | 0.0484 | 3.3e-37 | 0.136 |
| Fun with Fractals | 9.7 | 2.4e-21 | 0.61 | 0.0185 | 7.3e-22 | 0.079 |
| The Present | 13.5 | 2.0e-38 | 0.85 | 0.0396 | 8.8e-39 | 0.142 |

**Fig. 2.**
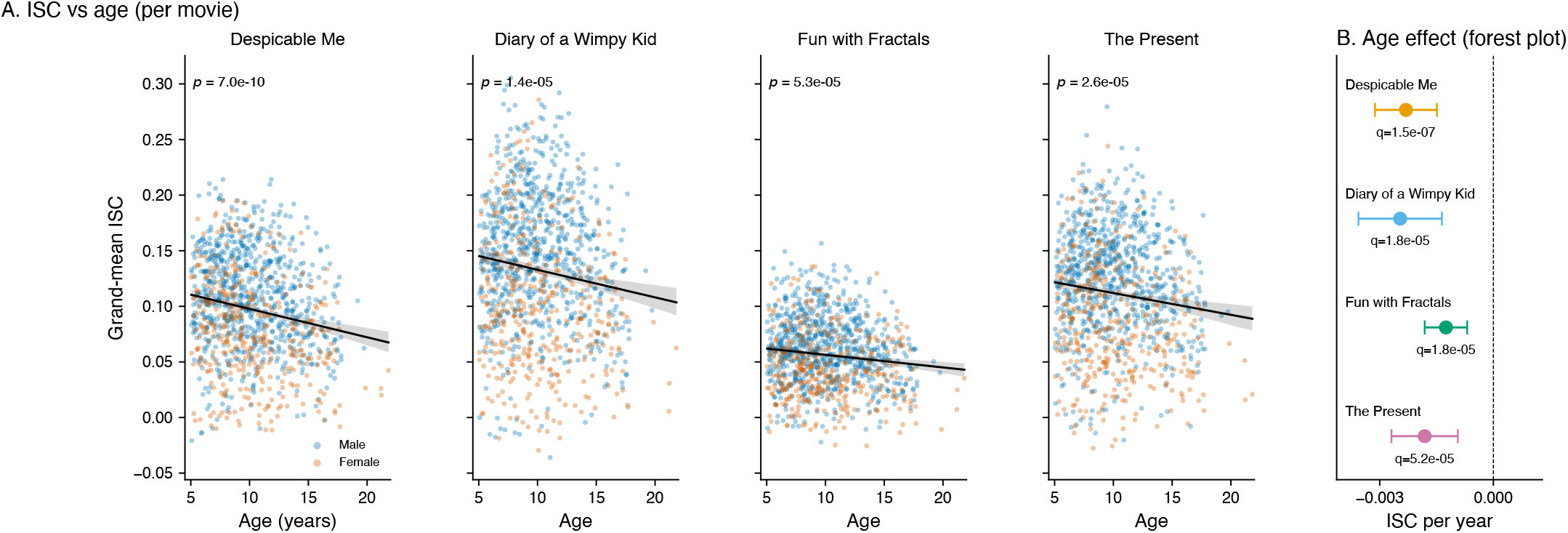
Developmental decline in EEG ISC replicates across all four movies. (A) Grand-mean ISC plotted against age for each movie (*n* = 1139 to 1143 each), shared y-axis across panels. Unadjusted linear fits with 95% CI shown in black; subjects colored by sex. All four slopes are negative at *p <* 1 *×* 10^−4^. (B) Forest plot of the age coefficient with 95% CI, in ISC per year, from the covariate-adjusted model with the quadratic term removed (sex, age, EHQ, release, four CBCL factors). All four are negative and exclude zero, in the same direction as the unadjusted fits in panel A.

The trajectory is not exactly linear. Retaining the quadratic term, the age^2^ coefficient is negative in all four movies (*p* = 1.0 *×* 10^−3^ to 8.9 *×* 10^−9^) with the implied vertex at 7.8 to 9.6 years, so ISC is roughly flat from age 5 to about 9 and then falls, with the decline steepening through adolescence. Three of the four movies show their highest raw bin mean at 8 to 11 rather than 5 to 8, by 0.9 to 3.3 percent, a rise an order of magnitude smaller than the subsequent fall. The net direction across the range tested is a decline, replicating Petroni’s finding of decreasing ISC, equivalently increasing neural variability, with age; their two-group design could not resolve this shape. We characterize the developmental pattern of the sex-difference component of ISC separately in Section 3.4.7, where the peak falls later, at 11 to 14 years.

This replication serves as a positive control for the analytic pipeline: a published developmental ISC effect from a smaller-cohort study (*n* = 114 + *n* = 303 in Petroni’s two cohorts) was recovered at *q <* 0.001 across all four movies in the present cohort (*n* = 1143), supporting confidence in the analytic procedures used to test the primary sex hypothesis below.

### 3.4 Sex Differences in ISC

#### 3.4.1 Primary Sex Effect

Males showed systematically higher ISC than females across all four movies. In unadjusted two-sample t-tests, sex effect t-statistics ranged from 9.68 (Fun with Fractals) to 13.46 (The Present), all with *p <* 3 *×* 10^−21^ (Table 4, Figure 3A). The magnitude of the effect, measured as the difference in grand-mean ISC between male and female subjects, ranged from 0.018 (Fun with Fractals) to 0.049 (Diary of a Wimpy Kid).

**Fig. 3.**
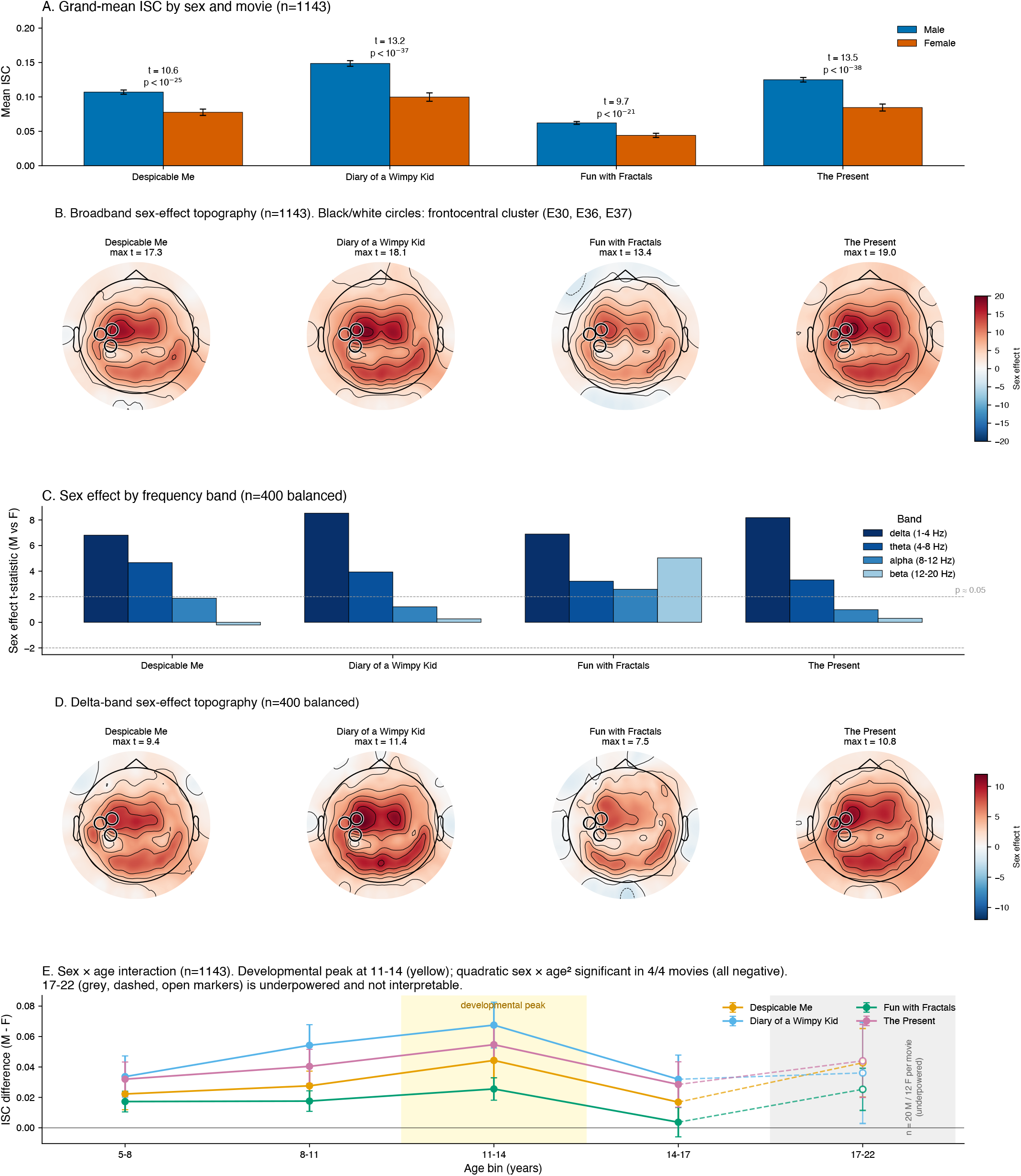
Robust low-frequency frontocentral male advantage in EEG ISC, with the overall sex difference peaking at ages 11 to 14. (A) Grand-mean ISC by sex across the four movies (*n* = 765 M, 378 F). All four show males higher than females at *p <* 3 *×* 10^−21^ (unadjusted t-tests). (B) Per-channel sex effect t-statistics on broadband ISC. Black/white circles mark the frontocentral cluster (E30, E36, E37) which is the t-statistic peak across all four movies, despite the raw effect spanning 82 to 100% of channels. (C) Sex effect t-statistics by frequency band (delta 1 to 4 Hz, theta 4 to 8 Hz, alpha 8 to 12 Hz, beta 12 to 20 Hz; *n* = 400 balanced subsample). The effect is largest in delta and present in theta across all four movies, and is negligible in alpha and beta except in the abstract control. (D) Delta-band sex effect topography on the *n* = 400 subsample. The delta map tracks the broadband map closely (spatial correlation *r* = 0.86 to 0.98 across movies), indicating that the broadband sex effect is largely the delta effect, though it is spatially broader than the theta effect and less tightly confined to the three-channel cluster. (E) Male-female ISC difference by age bin and movie. Among the interpretable bins the difference peaks at ages 11 to 14 (yellow shading) in all four movies, and the inverted-U is confirmed by significant negative quadratic sex *×* age^2^ interactions in 4 of 4 movies (all *q*_FDR_ < 0.05). The 17 to 22 bin (gray shading, dashed connector, open markers) contains 20 male and 12 female subjects in every movie and is not interpretable; its confidence intervals are the widest of the five bins but still understate its instability, since ISC in that range is both lower and less variable.

In the primary regression model with all demographic and clinical controls, the `sex_M` coefficient remained significant at *p <* 0.001 in all four movies, with coefficients ranging from +0.0185 (Fun with Fractals) to +0.0484 (Diary of a Wimpy Kid). Effect sizes were medium-to-large by Cohen’s conventions (unadjusted *d* = 0.61 to 0.85 across movies); partial *η*^2^ for `sex_M` after all controls was 0.08 to 0.14, making sex the largest single contributor to ISC variance among modeled covariates. Models explained 14% to 24% of variance in grand-mean ISC.

The direction of this effect is the same as that reported by Petroni et al. [9], who found higher ISC in male than female children in both of their cohorts. In their data the effect was marginal once age was partitioned: *F* (1, 87) = 3.83, *p* = 0.05 in the main cohort of 114, and *F* (1, 291) = 2.59, *p* = 0.1 in the replication cohort of 303, with the difference present among younger participants (*t*(53) = 2.02, *p* = 0.05 and *t*(224) = 2.29, *p* = 0.02 respectively) and absent among older ones. The present cohort recovers the same effect at *t* = 9.7 to 13.5 under a different ISC estimator. Sections 3.4.3 through 3.4.7 characterize its structure, and Section 3.4.4 addresses the alternative account raised in the original report.

#### 3.4.2 Robustness to Demographic and Clinical Confounding

Four robustness analyses were conducted. Three test whether the sex effect was driven by demographic or clinical compositional differences between male and female subgroups in the HBN sample; the fourth addresses partial cohort overlap with prior work. Two were prespecified (age stratification, strict QC subset) and are reported regardless of outcome. Two were added post-hoc: a low-symptom subset addressing clinical-sample confounding, and a release-exclusion analysis. Males in the cohort showed elevated attention CBCL scores compared to females (*M* = 0.12, *F* = −0.13, *p* = 2 *×* 10^−6^), consistent with the male preponderance of ADHD-spectrum presentations in the HBN referred sample; all four CBCL factors are entered as covariates in the primary regression.

##### Age stratification

When the sex effect was tested separately within each of five age bins, all 16 movie *×* age-bin cells (excluding the sparse 17 to 22 cell) showed the male advantage in the same direction; 15 of 16 cells reached raw *p <* 0.05. The effect was largest in the 11 to 14 age bin, suggesting a developmental peak rather than a stable trait difference. We return to this in Section 3.4.7.

##### Strict QC subset

Restricting the analysis to subjects passing strict QC criteria (*n* = 486) preserved the sex effect at *p <* 0.05 in all four movies, with effect sizes about 27% smaller (Cohen’s *d*) and correspondingly lower test statistics (*t* = 4.5 to 5.9). This effect-size reduction is genuine rather than sampling noise: the strict-QC *d* fell below 97 to 100% of 2000 random draws matched on sample size from the full cohort. Because *d* is normalized for sample size, the reduction is not a power effect, and its source is differential range restriction driven by a sex difference in data quality. Female recordings were on average slightly noisier than male (16.6 vs 15.2 *µ*V, *p* = 0.008), and because the absolute noise threshold removes the noisiest recordings, it retained proportionally fewer females (36% vs 46%), preferentially those with the lowest ISC; female mean ISC therefore rose more than male under strict QC and the sex gap compressed. A symmetric control that removed the same fraction of the noisiest recordings within each sex separately cost only 6% of *d*, versus 27% for the sex-blind threshold, indicating that roughly four fifths of the shrinkage reflects this asymmetry in what strict QC removes rather than loss of the underlying effect. Consistent with this, strict QC removed noise rather than signal: the sex effect was as large in the cleanest recording tertile (*d* = 0.83) as in the noisiest (*d* = 0.86), and mean ISC increased under strict QC.

##### Low-symptom subset (post-hoc)

Because the HBN cohort is recruited on clinical concern and lacks DSM diagnostic codes in the public BIDS release, we defined a low-symptom subset as subjects with all four CBCL bifactor scores below 0.5 SD above the cohort mean (*n* = 312, 27.3% of the analytic cohort; *n*_*M*_ = 195, *n*_*F*_ = 117). Although this subset cannot be equated with a population-level typical-development sample, it represents the subset of HBN subjects with the lowest parent-reported symptom burden across all four bifactor dimensions, providing the closest available approximation to a low-clinical-load comparison group within the public release. The sex effect persisted across all four movies (Despicable Me *t* = 4.11, *p* = 5.0 *×* 10^−5^, *d* = 0.48; Diary of a Wimpy Kid *t* = 6.45, *p* = 4.2 *×* 10^−10^, *d* = 0.76; Fun with Fractals *t* = 4.61, *p* = 6.0 *×* 10^−6^, *d* = 0.54; The Present *t* = 5.65, *p* = 3.5 *×* 10^−8^, *d* = 0.66). The frontocentral t-statistic topography also reproduced cleanly in this subset. Because diagnostic codes are not available in the public BIDS release, we cannot rule out residual confounding from undiagnosed clinical traits; the low-symptom replication therefore narrows but does not eliminate the clinical-sample confound.

##### Release exclusion (post-hoc)

The replication cohort of Petroni et al. [9] was drawn from HBN, so the present cohort partially overlaps theirs. Subject identifiers for their sample are not published, so we cannot identify the overlap directly and instead excluded conservatively by release, on the basis that their sample predates the later releases. Excluding R1 (*n* = 1026 subjects retained), R1 and R2 (*n* = 912), and R1 through R3 (*n* = 774) left the effect intact in every movie at every exclusion level, with all *p <* 1 × 10^−20^ (Supplementary Table S9). Coefficients did not decrease under exclusion. The effect is therefore not an artifact of the overlapping subjects; we do not interpret that direction as evidence in its own right, since release composition differs on several dimensions and release is already modeled as a fixed effect.

##### Full regression with all controls

A regression including sex, age, age^2^, EHQ, release, and all four CBCL factors as simultaneous predictors yielded `sex_M` coefficients significant at *p <* 0.001 in all four movies. Notably, none of the four CBCL factors were significant in any movie at *p <* 0.05 in the same regression, confirming that the sex effect is not mediated by the dimensional psychopathology measures available in HBN.

The sex effect was preserved across all four robustness analyses and the full regression, narrowing the role of demographic and clinical confounding as explanations for the observed pattern.

#### 3.4.4 Robustness to Template Composition, Ocular Artifacts, and Channel Interpolation

Three analyses tested whether the effect could arise from the measurement, or from the structure of the ISC computation itself, rather than from neural activity. The third, on channel interpolation, was added post-hoc. A fourth analysis, on evoked response magnitude, is reported separately in Section 3.4.4. Full numerical results are reported in Supplementary Materials.

##### Same-sex and size-matched templates

Because the leave-one-out template is the mean of all other subjects, the 2:1 male majority makes that template male-weighted, which could inflate male ISC for a purely compositional reason. We therefore recomputed ISC with same-sex templates, correlating each subject only against others of the same sex. On the balanced *n* = 400 subsample, where the two same-sex templates are already size-matched, the theta fronto-central effect strengthened rather than weakened (Hedges *g* from 0.889 to 1.274). On the full cohort, where the male template pools more subjects and is therefore less noisy, we additionally subsampled the male template to the female sample size in each movie (378, and 377 for The Present, which excludes the truncated recordings) and averaged over 50 random draws; the effect again strengthened (fronto-central cluster *g* from 1.376 to 1.751; whole-head grand mean *g* from 0.873 to 1.140). The response to the manipulation was asymmetric: reducing both templates by an identical amount left male ISC essentially unchanged while lowering female ISC by approximately 40 percent. Some asymmetry is expected on noise grounds alone, because a group with lower baseline ISC is more noise-limited and loses proportionally more when the template shrinks; the observed asymmetry is substantially larger than that mechanism predicts. The pooled mixed-sex template had, if anything, slightly understated the male advantage. Petroni et al. [9] computed their components within each sex group separately, so their sex comparison was likewise not subject to a mixed-template composition artifact.

##### Ocular artifact controls

Because ISC is sensitive to any stimulus-locked signal, including stimulus-locked eye movements, and because ocular artifacts project frontally, we tested whether the frontocentral effect could reflect residual ocular activity surviving ICA. The spatial evidence argues against an ocular origin: the sex-effect t-statistic peaks on the frontocentral cluster (electrodes E30, E36, E37; delta mean *t* = 9.2, theta mean *t* = 5.3) and is statistically null at the periocular electrodes nearest the eyes (delta mean *t* = 1.5, theta mean *t* = 0.6), the reverse of the periocular maximum that blink or saccade residual would produce, while the higher bands that would carry broadband saccadic-spike artifact (alpha, beta) show no sex effect except in the abstract control. The pattern is strongest in delta (1 to 4 Hz), the band in which any surviving blink energy would be most concentrated, which is the opposite of what an ocular origin predicts. We then tested the question directly by re-fitting the ICA per subject on a balanced subsample (*n* = 150, 75 male and 75 female; 149 for The Present, which excludes one truncated recording) and recomputing the effect with the ocular components removed, retained, or left in place. The frontocentral effect was essentially invariant across all three conditions (Table 5), unchanged in delta (*g* = 1.13, 1.16, and 1.15) and varying by 16 percent in the weaker theta effect (*g* = 0.75, 0.74, and 0.65) without approaching zero, whereas the same manipulation moved delta ISC by 24.4 percent at the ten periocular electrodes nearest the orbits (E1, E8, E14, E17, E21, E25, E32, E125, E126, E127) but by only 1.5 percent at the frontocentral cluster, a 16-fold dissociation pooled across movies, 9.9 to 49.7 fold within movies, showing that the ocular components reach the eyes but not the cluster. The dissociation held in all four movies. In the subset of subjects with at least one eye blink component (*n* = 112), the isolated ocular signal carried no detectable frontocentral sex difference in either band (delta *g* = −0.06, *p* = 0.77; theta *g* = 0.01, *p* = 0.96), and the sex effect was undiminished after controlling for it. As a further check on movement and drift contamination, we correlated per-recording amplitude (median per-channel standard deviation) with band-specific ISC. Delta ISC correlated negatively with amplitude (*r* = −0.26, *p <* 10^−6^), the only band showing a significant correlation, which is the opposite of what movement or drift contamination would produce. Petroni et al. [9] addressed ocular artifact by regressing out simultaneously recorded electrooculogram channels rather than by component removal; the two approaches converge on the same direction of effect.

**TABLE 5.**
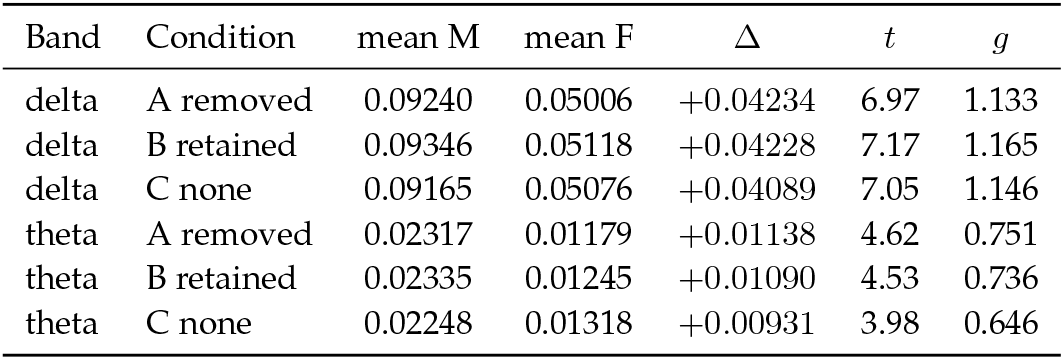
Frontocentral cluster sex effect under three ocular-removal conditions, from the balanced *n* = 150 subsample (75 male, 75 female). A: ocular components removed, the published condition. B: ocular components retained, all other artifact components removed. C: nothing removed. All three conditions derive from a single per-subject independent component decomposition, so they differ only in the exclusion list. The effect is unchanged across conditions in delta and varies by 16 percent in theta, in neither case approaching zero.

##### Channel interpolation (post-hoc)

Channels flagged as bad were spherically interpolated from their neighbors (Section 2.4), so a sex difference in how often channels are interpolated could in principle bias ISC. Female recordings carried more interpolated channels than male recordings in all four movies (pooled 23.91 against 21.86 of 129 channels, Welch *t* = 7.00, *p <* 0.001; per-movie differences of 1.47 to 2.73 channels, all *p ≤* 0.011). Interpolation count correlated negatively with grand-mean ISC in every movie (*r* = − 0.29 to − 0.36, all *p <* 1 *×* 10^−23^), indicating that in these data the count acts as a proxy for recording noise and that the associated noise penalty outweighs any inflation from spatial smoothing. The asymmetry therefore depresses female ISC and widens rather than narrows the observed gap.

Because the average reference precedes detection (Section 2.4), the recovered Cz channel is itself screened, and it is flagged far more often in female than in male recordings (35.45 against 6.33 percent in the *n* = 150 subsample, Fisher *p* = 1.3 *×* 10^−19^), consistent with its post-reference trace being the negative mean of all other channels and therefore a readout of montage-wide noise. The sex difference in flagged channels is not created by this mechanism: among recordings in which Cz was never flagged it remains 22 against 19 channels (Mann-Whitney *p* = 3.9 *×* 10^−3^). Excluding Cz from the 129-channel grand mean changes the four primary sex coefficients by 0.26 to 0.69 percent, in every case a reduction.

Entering interpolation count into the primary regression as an additional covariate reduced the `sex_M` coefficient by approximately 10 percent in every movie, retaining 87.7 to 90.8 percent of its magnitude (adjusted *β* = +0.0167 to +0.0425, all *p <* 2 *×* 10^−19^; partial *η*^2^ 0.071 to 0.126, against 0.079 to 0.142 without the covariate). The covariate itself was negative and significant in every movie (*β* = − 0.00095 to − 0.00202, all *p <* 1 *×* 10^−21^). The sex effect is therefore attenuated but not explained by differential interpolation.

The asymmetry is spatially concentrated at the fronto-central cluster. In the balanced *n* = 150 subsample for which per-channel interpolation records were retained, at least one of E30, E36, and E37 was interpolated in 36.1 percent of female recordings against 11.3 percent of male recordings (Fisher exact *p <* 0.001), a ratio of 3.2, whereas the whole-head interpolation ratio in the same subsample is 1.2 (24.39 against 20.37 channels). Interpolation at the cluster lowers cluster delta ISC substantially (0.042 against 0.080, a reduction of 47.8 percent). Restricted to recordings in which no cluster channel was interpolated, the fronto-central male advantage persists (delta Hedges *g* = 0.585, *p* = 8.5 *×* 10^−10^; theta *g* = 0.341, *p* = 3.6 *×* 10^−4^; *n* = 266 male, 191 female) against *g* = 0.975 in the interpolated stratum, which is heavily imbalanced at 34 male against 108 female recordings, and *g* = 0.760 pooled across both, all three computed at the recording level, an attenuation of 23 percent. The value of 1.133 in Table 5 is computed at the subject level and is not the comparator for these figures. The clean stratum is defined by cluster interpolation but selects on recording quality more generally, and two controls separate those. Removing recordings at the same sex-specific rates but at random reproduces none of the attenuation (mean *g* = 0.756 over 200 draws from the same subjects, 95 percent interval 0.677 to 0.852), so it is not a consequence of asymmetric sample loss alone. Removing them at the same rates ranked on interpolation outside the cluster gives *g* = 0.655, between the pooled and clean values, but with a bootstrap interval of 0.411 to 0.916 spanning both, from a cluster bootstrap resampling subjects with the statistic computed at the recording level, so the share attributable to cluster interpolation specifically is not resolvable at this subsample size. The same limit applies to the region comparison: the clean-to-pooled ratio is 0.770 at the cluster (95 percent CI 0.582 to 0.911) against 0.599 for the whole-head delta effect (0.203 to 0.865), with the interval on their difference including zero (− 0.486 to +0.030). Both ratios exclude zero, so a male advantage survives in the clean stratum in both regions. The asymmetry does not appear to place the topographic maximum at the cluster, on a weaker evidential footing than the effect-size analyses above. Recomputing the per-channel sex-effect map within the clean stratum leaves E30 the maximum-t channel in both delta and broadband, with spatial correlation 0.98 against the unstratified map over the same 599 recordings, whereas removing the same numbers of recordings ranked on interpolation outside the cluster shifts the maximum to E29 in delta and E36 in broadband. The identity of the maximum is unstable to the removal of 142 recordings by any rule at this subsample size: under random removal at the same sex-specific rates the maximum falls at E29 in 84 of 200 delta draws and at E30 in 77 of 200, and at E30 in 127 of 200 broadband draws. Both delta outcomes therefore lie within the range random removal produces, so the contrast between them is not itself evidence. What the analysis supports is the weaker statement that removing the recordings the asymmetry affects does not move the maximum away from the cluster, which an account on which the asymmetry places it there would predict. It does not establish that the location is independent of recording quality in general. This analysis is computed on the primary preprocessing pipeline rather than on the per-subject refit used for the ocular conditions of Table 5, which is also the basis of the stratum effect sizes above. On the same recordings and the same stratification the primary pipeline gives larger effect sizes in every stratum (clean *g* = 0.638, interpolated 1.167, pooled 0.838), with the same ordering and a clean-to-pooled ratio of 0.762 against 0.770, so the two bases agree on the structure and differ on the level. Two limits bound this analysis: per-channel interpolation records were retained only for the *n* = 150 subsample, so the cluster percentages carry that sampling uncertainty, and the regression covariate is a whole-head count used as a proxy for cluster-specific interpolation.

#### 3.4.4 Evoked Response Magnitude

Petroni et al. [9] reported that the sex effect on ISC did not survive control for steady-state visual evoked potential magnitude, and raised this as a candidate account under which the effect would reflect stronger stimulus-evoked responses in males rather than more stereotyped ones. Because ISC is a correlation, this is a genuine alternative: a larger evoked response relative to fixed background activity raises the correlation without any change in how shared the response is. We reproduced their control directly using the HBN Surround Suppression task, which employs the same paradigm at the same flicker frequency (Section 2.9).

##### Measure validity

SSVEP SNR peaked at electrode E76, with all five analysis electrodes (E76, E75, E71, E72, E77) posterior, consistent with a primary visual response (Figure 4A). SNR scaled with foreground contrast across the four levels (1.0008, 1.1620, 1.2444, 1.3056 at 0%, 30%, 60%, 100%), sitting at the noise floor in the no-flicker condition as expected. The paired within-subject excess of high-contrast (at or above 60 percent) over zero-contrast SNR was +0.2743 (SD 0.3569, *t*(353) = 14.46, Cohen’s *d* = 0.77), positive in 85.0 percent of subjects. Split-half reliability across odd and even trials was *r* = 0.891, or 0.943 after Spearman-Brown correction.

**Fig. 4.**
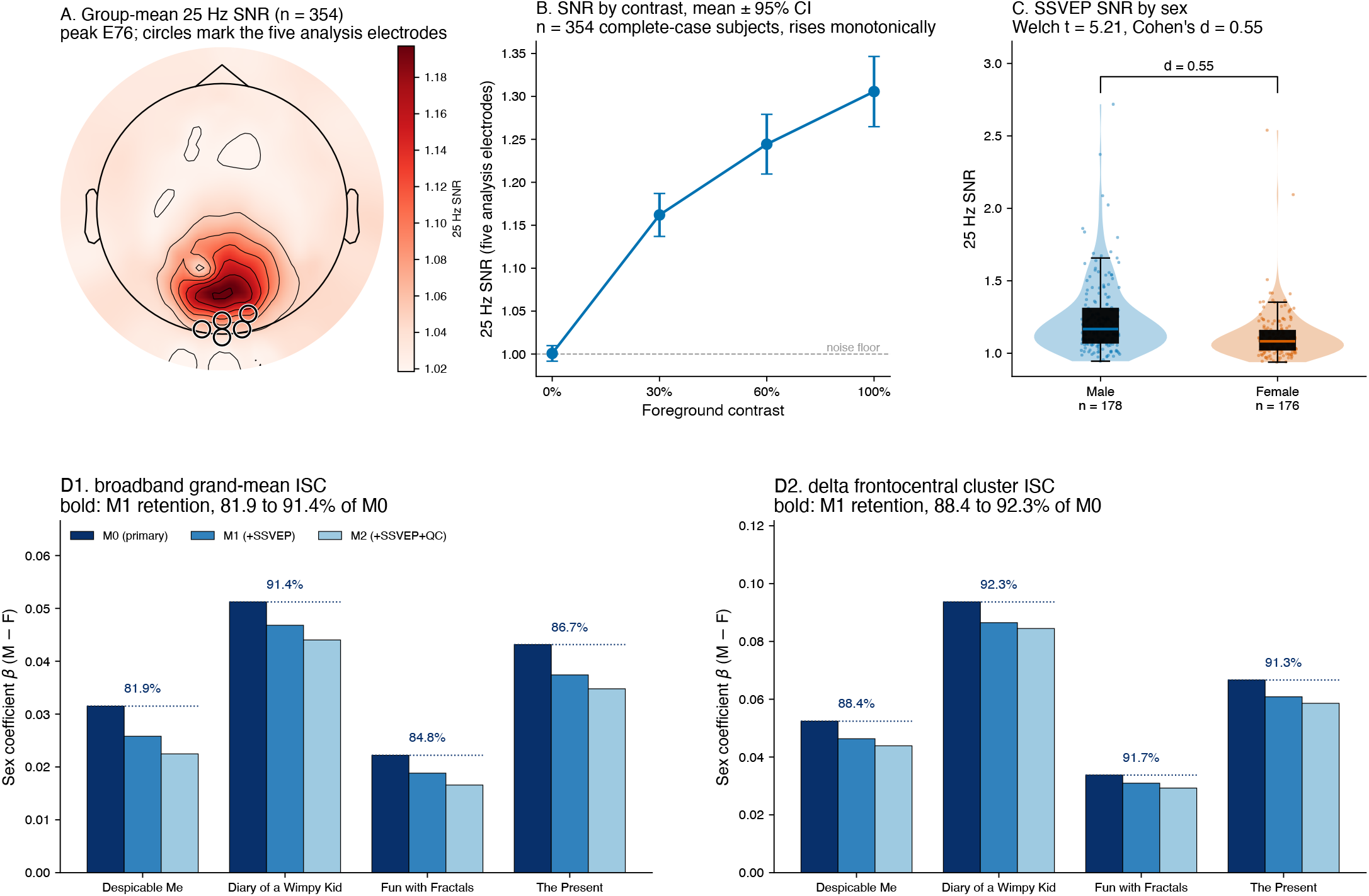
Evoked response magnitude does not account for the sex effect. (A) Group-mean 25 Hz signal-to-noise ratio across the scalp on the Surround Suppression task (*n* = 354), peaking at posterior electrodes as expected for a primary visual response. Analysis electrodes marked. (B) SSVEP SNR by foreground contrast, mean *±*95% CI, sitting at the noise floor with no flicker and rising monotonically with contrast. (C) SSVEP SNR by sex; males exceed females at *d* = 0.55. (D) Sex effect coefficient on ISC under M0, M1 and M2 for broadband grand-mean ISC and for delta frontocentral cluster ISC, per movie, showing retention of 81.9 to 91.4 percent on broadband and 88.4 to 92.3 percent on the delta cluster under M1.

##### Sample

Of the balanced *n* = 400 subsample, 354 have Surround Suppression recordings in the public release; extraction excluded no further subjects. Availability was balanced by sex (178 male and 176 female with recordings, 22 male and 24 female without; Fisher exact *p* = 0.876) and did not differ by age, general psychopathology factor, attention, or recording noise (all *p >* 0.32). Subjects without recordings had slightly lower grand-mean ISC (0.0774 against 0.0940, *t* = 2.49, *p* = 0.015).

##### Sex difference in SSVEP

SSVEP SNR was higher in male than female participants (male mean 1.2377, SD 0.2552, *n* = 178; female mean 1.1174, SD 0.1721, *n* = 176; Welch *t* = 5.21, *p* = 3.5 *×* 10^−7^, Cohen’s *d* = 0.55). The two coherently averaged measures gave the same result (*d* = 0.60 and *d* = 0.57). This difference did not reach significance in the original report (*F* (1, 106) = 3.3, *p* = 0.08), so the premise of the mechanism proposed there, that evoked magnitude differs by sex, is supported at this sample size. Whether it accounts for the ISC effect is addressed below.

##### Correlation with ISC

SSVEP SNR correlated positively with grand-mean ISC in every movie (*r* = 0.202 to 0.251, all *p <* 2 × 10^−4^), against *r* = 0.41 at *N* = 84 in the original report.

##### Effect of control

Adding SSVEP SNR to the primary regression left the sex effect significant in 4 of 4 movies for both outcomes (Table 6, Figure 4D). On broadband grand-mean ISC the coefficient retained 81.9 to 91.4 percent of its magnitude, and on delta frontocentral cluster ISC it retained 88.4 to 92.3 percent, with all *p <* 8 *×* 10^−11^. Adding quality-control covariates alongside SSVEP retained 71.3 to 86.0 percent and 83.8 to 90.2 percent respectively. The two coherently averaged SSVEP measures gave retention of 78.1 to 92.3 percent on broadband.

**TABLE 6.** Sex effect on ISC before and after control for evoked response magnitude, on the *n* = 354 subsample with Surround Suppression recordings. M0 is the primary regression; M1 adds SSVEP signal-to-noise ratio at 25 Hz; M2 adds quality-control covariates alongside SSVEP. Retained is the M1 or M2 coefficient as a percentage of M0. Mediation is the proportion of the M0 coefficient carried by the path through SSVEP, with a bootstrap 95% confidence interval on the indirect path (2000 resamples). The upper block is broadband grand-mean ISC; the lower block is delta-band ISC at the frontocentral cluster (E30, E36, E37). All M1 and M2 coefficients significant at *p <* 1.6 *×* 10^−5^; all delta cluster M1 and M2 at *p <* 8 *×* 10^−11^. Regression *n* = 353 per movie and 352 for The Present, against the 354 subjects with usable SSVEP extractions: one subject is removed by the listwise deletion of Section 2.7 and one further recording by the truncation exclusion of Section 2.1.

| Movie | M0 |  | M1 (+SSVEP) |  | M2 (+SSVEP+QC) |  | Mediation |  |
| --- | --- | --- | --- | --- | --- | --- | --- | --- |
| | $\beta$ | $t$ | $\beta$ | retained | $\beta$ | retained | % | 95% CI |
| *Broadband grand-mean ISC* |  |  |  |  |  |  |  |  |
| Despicable Me | 0.0315 | 6.13 | 0.0258 | 81.9% | 0.0225 | 71.3% | 18.1 | 0.0027, 0.0099 |
| Diary of a Wimpy Kid | 0.0512 | 8.21 | 0.0468 | 91.4% | 0.0440 | 86.0% | 8.6 | 0.0011, 0.0093 |
| Fun with Fractals | 0.0222 | 6.85 | 0.0188 | 84.8% | 0.0166 | 74.7% | 15.2 | 0.0015, 0.0059 |
| The Present | 0.0432 | 8.10 | 0.0374 | 86.7% | 0.0348 | 80.6% | 13.3 | 0.0028, 0.0096 |
| *Delta frontocentral cluster ISC* |  |  |  |  |  |  |  |  |
| Despicable Me | 0.0524 | 8.57 | 0.0463 | 88.4% | 0.0439 | 83.8% | 11.6 | 0.0017, 0.0104 |
| Diary of a Wimpy Kid | 0.0937 | 11.53 | 0.0865 | 92.3% | 0.0845 | 90.2% | 7.7 | 0.0025, 0.0136 |
| Fun with Fractals | 0.0338 | 8.06 | 0.0310 | 91.7% | 0.0293 | 86.7% | 8.3 | 0.0007, 0.0053 |
| The Present | 0.0667 | 10.65 | 0.0608 | 91.3% | 0.0586 | 87.9% | 8.7 | 0.0022, 0.0102 |

All M1 and M2 coefficients significant at $p < 1.6 \times 10^{-5}$ ; all delta cluster M1 and M2 at $p < 8 \times 10^{-11}$ . Regression $n = 353$ per movie and 352 for The Present, against the 354 subjects with usable SSVEP extractions: one subject is removed by the listwise deletion of Section 2.7 and one further recording by the truncation exclusion of Section 2.1.

##### Mediation

The mediated proportion is the complement of the retention above rather than an independent quantity: 8.6 to 18.1 percent of the broadband sex effect runs through the path via SSVEP magnitude, and 7.7 to 11.6 percent of the delta frontocentral effect. What the decomposition adds is an interval: all bootstrap confidence intervals on the indirect path exclude zero (Table 6), so the path is non-zero as well as small. The mediated share is smaller for the frontocentral delta outcome than for the whole-head broadband outcome in all four movies.

Because SSVEP magnitude itself differs by sex, it is a mediator of the sex effect rather than a confound of it, and conditioning on a mediator biases the direct effect downward. The retained coefficients should therefore be read as conservative.

#### 3.4.5 Topography of the Sex Effect

The sex effect was near-global in spatial extent. Per-channel t-tests with FDR correction across 129 channels within each movie revealed that 82% to 100% of channels showed significant male *>* female ISC, with 0 to 1 channels per movie showing the opposite direction (Figure 3B, Table 7). However, the t-statistic peak was localized to a frontocentral core cluster of E30, E36, and E37, with adjacent E31 in the top five in three of the four movies and E29 in two. E30 was the maximum-t channel for all four movies in broadband. These channels lie over central / left frontocentral cortex in the GSN-HydroCel-129 layout, approximately corresponding to FC1-FC3-F1 in the standard 10-20 system. In subsequent analyses we use “frontocentral cluster” to refer to the core three channels (E30, E36, E37) unless otherwise noted.

**TABLE 7.** Topographic distribution of the sex effect on per-channel ISC per movie. Top channels by sex effect t-statistic, with the count of channels surviving FDR correction at *q <* 0.05 in each direction. The frontocentral cluster (E30, E36, E37) consistently appears among top-5 channels across all four movies, with E30 the maximum-t channel in every movie.

| Movie | Top<br>5 channels | Max<br>$t$ | FDR +<br>/129 | FDR –<br>/129 |
| --- | --- | --- | --- | --- |
| Despicable Me | E30, E31, E36, E37, Cz | 17.3 | 120 | 0 |
| Diary of a Wimpy Kid | E30, E31, E36, E37, E80 | 18.1 | 127 | 0 |
| Fun with Fractals | E30, E37, E36, E29, E31 | 13.4 | 106 | 1 |
| The Present | E30, E36, E29, E105, E37 | 19.0 | 129 | 0 |

Two distinct topographic signatures emerged from this analysis: the raw male-female ISC difference was largest at parieto-occipital electrodes (where overall ISC is also largest), with spatial correlation between the difference map and the overall ISC topography of *r* = 0.58 to 0.67 across movies. In contrast, the t-statistic map peaked at the frontocentral cluster (spatial correlation with overall ISC topography: *r* = 0.27 to 0.45). The discrepancy reflects variance scaling: at parieto-occipital channels both sexes have relatively high ISC, reducing the proportional contrast despite a large raw difference; at frontocentral channels female ISC is small in absolute terms, magnifying the male advantage relative to noise.

#### 3.4.6 Frequency-Band Specificity

To localize the sex effect to specific neural frequency bands, ISC was recomputed in four bands (delta 1 to 4 Hz, theta 4 to 8 Hz, alpha 8 to 12 Hz, beta 12 to 20 Hz) on a balanced subsample of *n* = 400 (200 male, 200 female; sampled within sex with seed 42), using the same two-sample t-test and primary regression framework as the grand-mean analysis.

The sex effect was concentrated in the low-frequency bands and was largest in delta (Figure 3C, Table 8). Delta carried the strongest sex effect of any band, significant across all four movies in two-sample t-tests (*t* = 6.81 to 8.53, all *p <* 4 *×* 10^−11^) and in full regression with all controls (*p <* 3 *×* 10^−10^ in 4 of 4 movies). Theta was the next strongest, also significant in all four movies (*t* = 3.21 to 4.66, all *p <* 1.5 *×* 10^−3^; full regression *p <* 1.2 *×* 10^−3^ in 4 of 4 movies). Delta t-statistics exceeded theta by factors of approximately 1.5 to 2.5 in every movie; at the frontocentral cluster the corresponding effect sizes were Hedges *g* = 1.35 for delta against 0.89 for theta (Supplementary Materials). These low-frequency bands also dominated absolute ISC magnitudes, indicating that the cross-subject shared signal during movie viewing is concentrated at low frequencies.

**TABLE 8.** Sex effect on band-specific grand-mean ISC (the mean across the 129 channels) per movie, from the balanced band subsample (*n* = 400; 398 for The Present, which excludes two truncated recordings), computed with the same two-sample t-test (M vs F) as the broadband grand-mean analysis. Cluster-level and pooled-across-movie band statistics are reported in Supplementary Materials. Delta (1 to 4 Hz) and theta (4 to 8 Hz) carry the sex effect across all four movies, with delta the strongest; both are also significant in 4 of 4 movies in the full regression with all controls. Alpha and beta are null except in the abstract control (Fun with Fractals).

| Movie | Delta (1 to 4 Hz) |  | Theta (4 to 8 Hz) |  | Alpha / Beta |  |
| --- | --- | --- | --- | --- | --- | --- |
| | $t$ | $p$ | $t$ | $p$ | $t_\alpha/t_\beta$ | $p_\alpha/p_\beta$ |
| Despicable Me | 6.81 | 4e-11 | 4.66 | 4e-6 | 1.88 / -0.21 | 0.06 / 0.84 |
| Diary of a Wimpy Kid | 8.53 | 3e-16 | 3.93 | 1e-4 | 1.21 / 0.27 | 0.23 / 0.79 |
| Fun with Fractals | 6.90 | 2e-11 | 3.21 | 1.4e-3 | 2.58 / 5.04 | 0.01 / 7e-7 |
| The Present | 8.18 | 4e-15 | 3.31 | 1.0e-3 | 0.99 / 0.31 | 0.32 / 0.76 |

**TABLE 9.**
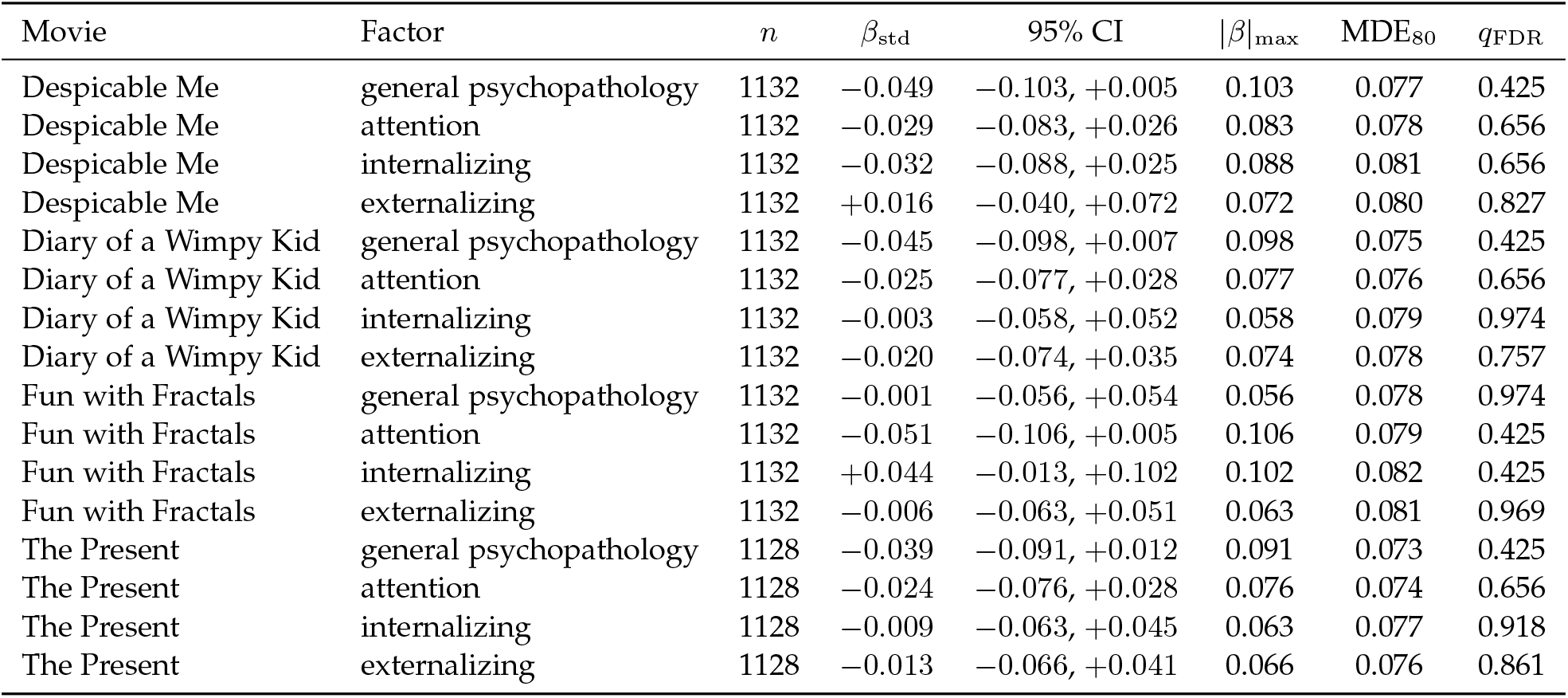
Bound on CBCL bifactor prediction from grand-mean ISC. Standardized regression coefficients with 95% confidence intervals, from models controlling for sex, age, age^2^, EHQ and release. |*β*| _max_ is the largest absolute association the confidence interval is consistent with; MDE_80_ is the minimum detectable effect at 80% power and two-sided *α* = 0.05 for the achieved standard error. FDR correction is across all 16 tests. No test reaches *q <* 0.05, and no confidence interval admits an association larger than 0.106.

Alpha and beta bands showed largely null sex effects in the regression with controls (alpha: 1 of 4 movies significant; beta: 1 of 4 movies significant; all surviving hits restricted to Fun with Fractals). We do not have an account of why Fun with Fractals alone shows sex differences in the higher bands. The beta effect in that stimulus (*t* = 5.04, *p* = 7 *×* 10^−7^) is larger than its theta effect, which no interpretation we have considered predicts. One possibility is that the abstract mathematical animation, lacking narrative or speech content, drives broader visual entrainment; we offer this only as a conjecture, since nothing in our data tests it. The result is reported as observed and left unexplained.

The delta effect shows the same frontocentral weighting and periocular null as the theta and broadband effects (delta mean *t* = 9.2 at the cluster against 1.5 at the periocular electrodes). Because delta is the band most exposed to ocular and movement artifact, we verified this topography and its robustness to ocular-component removal in detail (Section 3.4.3): the delta effect is frontocentral rather than periocular and survives removal of the ocular components, arguing against an artifactual origin. Full band results are reported in Supplementary Materials.

Per-channel sex effects were recomputed in the delta and theta bands on the *n* = 400 subsample to establish how the band-limited topographies relate to the broadband result. The delta map (Figure 3D) tracks the broadband map closely, with spatial correlations of *r* = 0.86 to 0.98 across movies, indicating that the broadband sex effect is largely the delta effect. Its maximum-t channel falls on a core cluster channel in two of four movies (E30 for Diary of a Wimpy Kid, E36 for The Present) and on an adjacent channel in the other two (E29 for Fun with Fractals, Cz for Despicable Me). Between 3 and 5 of the five frontocentral cluster channels (E30, E36, E37, E29, E31) appear in the delta top-10 per movie.

The delta effect is therefore spatially broader and more vertex-weighted than the theta effect, which is the more tightly localized of the two: in theta the maximum-t channel is a core cluster channel in three of four movies (E30, E37 and E37 for Despicable Me, Diary of a Wimpy Kid and The Present respectively) and the adjacent E31 for Fun with Fractals, at least 4 of those five channels appear in the top-10 in every movie, and spatial correlation with broadband is lower at *r* = 0.64 to 0.75. We therefore describe the topography of the sex effect as frontocentrally weighted rather than confined to the three-channel cluster. The cluster’s status as the t-statistic peak rests on the full-cohort broadband analysis (Table 7), where E30 is the maximum-t channel in all four movies at *n* = 1143, and not on the band-limited subsample. One qualification applies to that claim. Because interpolation at the cluster depresses cluster ISC, and female recordings are interpolated there disproportionately (Section 3.4.3), the interpolation asymmetry inflates the male-minus-female difference at the cluster specifically and could in principle contribute to the peak being located there rather than only to its height. Within the *n* = 150 subsample for which per-channel interpolation records were retained, this is testable, and the peak does not move: E30 remains the maximum-t channel among recordings with no cluster channel interpolated, in both delta and broadband, while two rate-matched removal controls do move it (Section 3.4.3). Per-channel records were not retained for the full cohort, so the test rests on 457 recordings, and at that size the identity of the maximum is sensitive to which recordings are removed. The topography is accordingly read as fronto-centrally weighted, with the location of the maximum tested against the interpolation asymmetry at subsample scale and unchanged by it, though not verified against it at full-cohort scale.

#### 3.4.7 Developmental Peak

The age-stratified sex effect showed the male advantage was largest in the 11 to 14 age bin across all four movies. To test this developmental inverted-U pattern formally, regressions were fit per movie with both linear `sex_M` *×* age (age centered within each movie at that movie’s mean) and quadratic `sex_M` *×* age^2^ interaction terms entered jointly, plus all standard covariates. The quadratic interaction was significant in 4 of 4 movies (Despicable Me: *β* = − 0.00048, *p* = 0.022, *q*_FDR_ = 0.029; Diary of a Wimpy Kid: *β* = − 0.00114, *p <* 1 *×* 10^−4^, *q*_FDR_ < 1 *×* 10^−3^; Fun with Fractals: *β* = − 0.00029, *p* = 0.044, *q*_FDR_ = 0.044; The Present: *β* = − 0.00070, *p* = 0.0017, *q*_FDR_ = 0.003), with all coefficients negative, indicating the sex effect is concave (inverted-U) about the centering age (Figure 3E). All four survived Benjamini-Hochberg FDR correction across the four movies at *q <* 0.05, the weakest being Fun with Fractals at *q*_FDR_ = 0.044. The linear interaction was significant in 3 of 4 movies in the joint model (*p* = 0.033, 0.024, 0.833, 0.031), indicating the inverted-U peak is shifted slightly from cohort-mean age. For comparison, a linear-only model (without the quadratic term) shows a non-significant sex *×* age interaction in all four movies (*p* = 0.10 to 0.46), confirming that the inverted-U pattern obscures a net linear effect when the quadratic component is omitted.

Visual inspection of the per-bin sex differences confirmed the inverted-U pattern in all four movies: the male-female ISC difference grew from 5 to 8 to 8 to 11 to a peak at 11 to 14, then declined into 14 to 17 (with 17 to 22 too sparse to interpret reliably; *n*_*M*_ = 20, *n*_*F*_ = 12 across the cohort).

This trajectory is consistent with the age-stratified pattern reported by Petroni et al. [9], who found the sex difference present among younger participants and absent among older ones in both of their cohorts. Their design contrasted two broad age groups rather than fitting a continuous trajectory, so it could establish the presence of the difference in childhood and its absence later, but not the location of the peak.

#### 3.4.8 Stimulus Specificity

A sex *×* movie interaction test on long-format data was significant (*F* (3, 4507) = 22.88, *p* = 1 *×* 10^−14^; *R*^2^ gain over the main-effects model 0.0095), but reflects magnitude rather than direction: the effect is positive in all four stimuli, ranging 2.6-fold from Fun with Fractals (+0.0185) to Diary of a Wimpy Kid (+0.0484), with the narrative-containing movies larger than the abstract control and the sex effect scaling with overall ISC magnitude.

### 3.5 ISC Does Not Predict CBCL Bifactor Scores

We tested whether grand-mean ISC predicted any of the four CBCL bifactor dimensions (general psychopathology factor, attention, internalizing, externalizing) across the four movies (16 tests total) using regressions controlling for sex, age, age^2^, EHQ, and release. After FDR correction across the 16 tests, no test reached *q <* 0.05 (Figure 5A). The minimum raw p-value was 0.07 (Despicable Me *×* p factor and Fun with Fractals *×* attention), with the minimum *q*_FDR_ at 0.425.

**Fig. 5.**
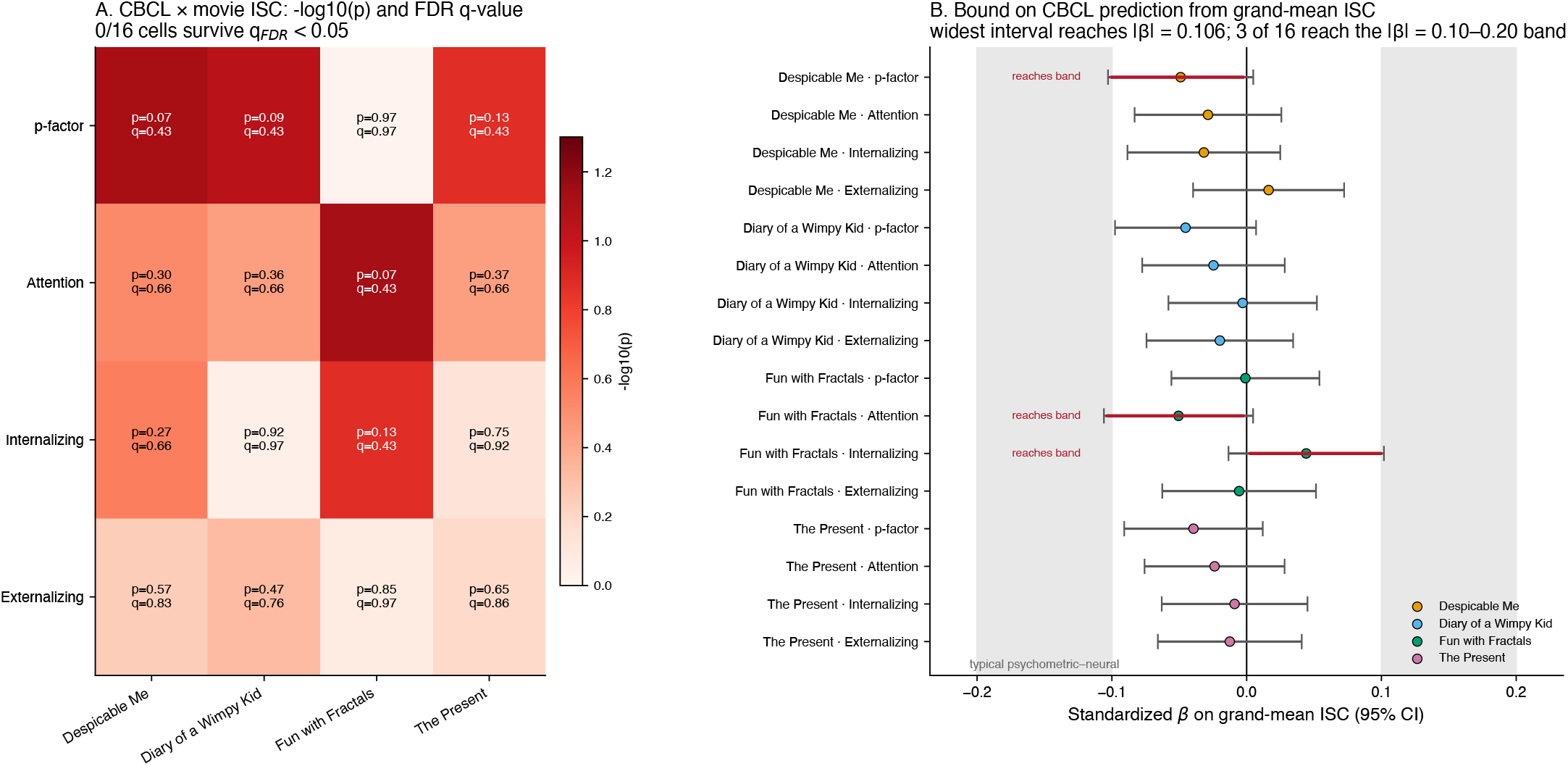
(A) ISC *×* CBCL bifactor regression results across 16 tests (4 movies *×* 4 factors), with sex, age, age^2^, EHQ, and release as covariates. No test reached *q*_FDR_ < 0.05; the minimum was 0.425. The null extends to per-channel ISC under joint FDR (0/2064 tests significant) and to the frontocentral cluster carrying the sex effect (0/16 tests significant). (B) Forest plot of the same 16 associations in standardized units with 95% confidence intervals. Shaded band marks the range of association magnitudes typically reported between psychometric instruments and neural measures (|*β*| = 0.10 to 0.20); three of the 16 intervals extend just into its lower edge and the remaining 13 fall short of it.

Because a null is uninformative without a bound, we report each association in standardized units with its 95% confidence interval (Table 9). The largest absolute standardized coefficient across all 16 tests was 0.051, and the widest confidence interval was consistent with no association larger than |*β*| = 0.106, corresponding to less than 1.2 percent of variance in grand-mean ISC. The median minimum detectable effect at 80 percent power was 0.078, so the design was powered to detect associations at the upper edge of what the confidence intervals permit. Associations of the magnitude typically reported between psychometric instruments and neural measures (*r* = 0.10 to 0.20) therefore lie above this bound except at its lower edge: three of the 16 intervals extend just past |*β*| = 0.10 (Despicable Me with general psychopathology, 0.103; Fun with Fractals with attention, 0.106; Fun with Fractals with internalizing, 0.102) and the remaining 13 fall short of it. Associations of 0.11 or larger are excluded at every test.

To verify that the grand-mean null was not an averaging artifact, we extended the analysis to the per-channel level. We ran 2064 regressions (4 movies *×* 4 CBCL factors *×* 129 channels) with the same covariate structure and applied joint Benjamini-Hochberg FDR across the full 2064-test family. No channel survived *q*_FDR_ < 0.05 in any movie *×* factor combination; the global minimum *q*_FDR_ across all 2064 tests was 0.92. The null does not depend on that family being the most stringent available. Correcting within each movie across the four factors and 129 channels (4 families of 516 tests) gives a global minimum *q*_FDR_ of 0.58, and correcting within each movie and factor separately (16 families of 129 tests) gives 0.20; zero tests survive under any of the three schemes. The smallest raw p-value among the 2064 per-channel tests is 0.0027, at one channel for The Present with the general psychopathology factor. We additionally tested mean ISC across the frontocentral cluster (E30, E36, E37) that carries the sex effect, against each of the four CBCL factors across the four movies (16 cluster-targeted tests); none survived *q*_FDR_ < 0.05 (minimum *q*_FDR_ = 0.91), and the bound at the cluster is tighter than at the grand mean, with the widest interval admitting no association beyond |*β*| = 0.086. The CBCL null is therefore not a property of the grand-mean summary statistic alone but extends to per-channel and cluster-targeted feature representations.

This null result is consistent with the absence of a CBCL *×* ISC effect at smaller scale (R1 to R4, 628 subjects, *n* = 619 after the listwise deletion of Section 2.7; minimum *q*_FDR_ = 0.82). The result indicates that ISC during pediatric naturalistic stimulus viewing, evaluated at multiple feature representations on the four-movie HBN protocol, does not encode CBCL-bifactor-dimensional psychopathology in this cohort at sample size and analytic resolution sufficient to detect typical-magnitude effects.

## 4 Discussion

### 4.1 Interpreting the Sex Difference in ISC

Male participants showed higher ISC than female participants in all four movies, an effect concentrated in the low-frequency bands and largest in delta, peaking in t-statistic at a frontocentral cluster and peaking developmentally at ages 11 to 14. Interpreting that pattern requires separating what the data show from what they mean. We first consider explanations under which the effect would not reflect a difference in shared neural processing, and then the interpretations that remain.

#### Explanations not requiring a difference in shared processing

Five are addressed by the analyses above, and each is either excluded or quantified. Template composition is not the source: the 2:1 male skew makes the leave-one-out template male-weighted, but same-sex and size-matched templates strengthened the effect rather than weakening it (Section 3.4.3). Ocular and movement artifact is not the source, a concern sharpened by the effect being largest in the band most exposed to it: the effect peaks frontocentrally rather than periocularly, is null in the higher bands that would carry saccadic-spike artifact except in the abstract control, is invariant to removal of the ocular components, and recording amplitude correlates negatively with delta ISC, the opposite of what drift contamination would produce. Differential noise is not the source: female recordings are slightly noisier, which compresses the effect under strict quality control (Section 3.4.2), but the effect is as large in the cleanest recordings as in the noisiest. Differential channel interpolation inflates but does not create it: adjusting for interpolation count costs roughly 10 percent, and the frontocentral advantage persists in recordings with no cluster channel interpolated, attenuated by 23 percent, of which the share attributable to cluster interpolation rather than to general recording quality is not resolvable at the available subsample size. Greater evoked response magnitude in males, the account raised in the original report, is real but insufficient: SSVEP magnitude does differ by sex and does predict ISC, yet carries 8.6 to 18.1 percent of the broadband effect and 7.7 to 11.6 percent of the frontocentral delta effect (Section 3.4.4).

The consistent picture is that several measurement-level and response-level asymmetries inflate the measured effect without accounting for it. These analyses narrow the non-neural explanations we are able to test and leave a difference in shared processing as the most plausible remaining account, but the interpretations below are hypotheses a single cohort cannot adjudicate.

#### Sample composition

HBN is community-referred on parent-reported concern, a design expected to yield a high proportion of participants affected by psychiatric illness, with ADHD prominent in the sample [20], and a roughly 2:1 male skew that follows from it. The male advantage might therefore reflect ADHD-spectrum traits influencing engagement with naturalistic stimuli, with boys engaging more strongly with rapid-cut content. This is partially testable here and is not strongly supported: attention CBCL is entered as a covariate in the primary regression, no CBCL factor predicted ISC in regressions including sex, the effect persists in the low-symptom subset, and the same directional effect appears in a cohort recruited independently of HBN [9], [10], which weakens any account resting on HBN’s referral structure. Residual confounding from unmeasured ADHD-related traits cannot be excluded, since CBCL scores are parent-reported and capture continuous variation rather than diagnostic specificity.

#### Maturational timing

Females typically show earlier maturation in language and social-cognitive networks during childhood [28]. If ISC is higher in less-mature brains, as proposed in the original report, the male advantage could reflect less individuated processing in male brains in this window. Our peak at 11 to 14 years sits where sex differences in cortical maturation are typically largest, and the original report likewise found the difference present in childhood and absent later. Our continuous trajectory adds that the difference rises to a pre-adolescent peak rather than declining monotonically from the youngest ages, which a simple maturational-lag account does not by itself predict.

#### Engagement and attention

Higher ISC tracks behavioral engagement, attention and memory in adult work, and the low-frequency frontocentral localization is suggestive here, since theta overlaps frequencies associated with attentional engagement and mentalizing. On this account male participants would sustain attention to narrative content more uniformly in this age range. The 2.6-fold difference in effect magnitude between Diary of a Wimpy Kid and Fun with Fractals is consistent with it, though the effect persists in the abstract control.

The most defensible reading is that the male advantage reflects some combination of these, and that adjudicating between them requires comparison cohorts not available in HBN.

### 4.2 Relation to Petroni et al. (2018)

Both of our principal developmental findings match theirs in direction: ISC declines with age, and among children male participants show higher ISC than female participants. Three features of their design bear on how the present results relate to them.

#### Effect strength

Their sex effect was strong pooled across stimuli (*F* (1, 393) = 53.11 in the main cohort; *F* (1, 823) = 11.12 in the replication) but marginal once age was partitioned: *F* (1, 87) = 3.83, *p* = 0.05, and *F* (1, 291) = 2.59, *p* = 0.1. Within the younger group it reached *t*(53) = 2.02, *p* = 0.05 and *t*(224) = 2.29, *p* = 0.02. We recover it at *t* = 9.7 to 13.5 in every stimulus, which is what allows the spectral, topographic and developmental structure reported here to be resolved.

#### Independence and overlap

Their two cohorts differ in a way that matters. The main cohort of 114 came from a separate dataset [10], so it constitutes an independent confirmation of the direction. The replication cohort of 303 came from HBN and therefore overlaps the present cohort to an unknown degree; subject identifiers are not published. Because their sample predates the later releases, we tested the effect with the earliest releases excluded (Section 3.4.2). Removing R1 through R3 discards 369 subjects, roughly a third of the cohort, and the effect remains at *p <* 1 *×* 10^−20^ in every movie. We do not interpret the direction of change in the coefficients, since release composition differs on several dimensions and release is already modeled as a fixed effect, but the analysis establishes that the effect does not depend on the overlapping subjects.

#### Methodological independence

The two studies estimate ISC differently. They used correlated component analysis on three maximally correlated components [5]; we average per-channel leave-one-out correlations across 129 channels. They removed ocular artifact by regressing simultaneously recorded electrooculogram channels; we removed classified ocular independent components. That two substantially different measurement chains yield the same direction is convergent evidence rather than redundancy. Consistent with this, the resting-state floor is effectively identical across the two studies (0.001 against 0.00133) and the rank ordering of ISC magnitude across the three shared stimuli is the same. The two estimators do not agree on effect magnitude, however. In their HBN replication cohort the age-ISC correlation was *r* = − 0.37 to − 0.44; over essentially the same age range in the same dataset we obtain *r* = − 0.12 to − 0.18 unadjusted, roughly a third as large despite a cohort nearly four times the size. Correlated component analysis fits its projections so as to maximize cross-subject correlation within the cohort it is applied to, so the recovered components carry whatever structure dominates that cohort, including its age composition, whereas a channel-space mean is fitted to nothing. We therefore read the direction and the ordering as replicating and treat magnitude comparisons between the two estimators as uninformative.

#### The evoked-magnitude question

The most substantive question they raised, and could not resolve, is whether the effect reflects shared processing at all. They found ISC correlated with SSVEP magnitude (*r* = 0.41, *N* = 84) and that the sex effect did not survive regressing SSVEP out (*F* (1, 81) = 0.08, *p* = 0.8), while noting the smaller sample. Our reproduction on 354 subjects supports the premise and not the conclusion. SSVEP magnitude does differ by sex, which their sample could not establish (*F* (1, 106) = 3.3, *p* = 0.08 there; *d* = 0.55 to 0.60, *p <* 4 *×* 10^−7^ here), and it does predict ISC. But the mediated path carries 8.6 to 18.1 percent of the broadband effect and 7.7 to 11.6 percent of the frontocentral delta effect, and the effect remains significant in every movie for both outcomes after control. Because SSVEP magnitude is itself sex-dependent, conditioning on it removes part of the effect of interest, so the retained coefficients are conservative.

A mediated share of this size does not by itself account for the reduction they observed. Their effect corresponds to approximately *d* = 0.42 before control and *d* = 0.06 after, a reduction of about 85 percent, whereas mediation of the size measured here predicts a residual of *d* = 0.34 to 0.38, corresponding to *F* (1, 81) near 2.4 to 3.0 and *p* near 0.09 to 0.13: still marginal at their sample size, but not absent. The remainder is most plausibly sampling variability acting on an already-marginal effect in a subsample reduced from 114 to 84. We therefore read their result as an underpowered test of a real mechanism rather than as a demonstration that the mechanism accounts for the effect.

### 4.3 Low-Frequency Frontocentral Specificity

Two convergent findings sharpen the neural interpretation: the effect is concentrated in the low-frequency bands, largest in delta and present in theta, and peaks at frontocentral electrodes (E30, E36, E37, approximately FC1-FC3-F1 in the standard 10-20 system).

Frontocentral theta is associated with sustained attention, mentalizing, error monitoring and the integration of narrative content over time [29], and theta synchronization has been proposed as a coordinating signal for the distributed networks that naturalistic stimuli engage [30]. That the effect is in fact largest in delta is consistent with slow cortical activity carrying much of the reliable stimulus-locked signal during continuous viewing, though the functional interpretation of delta during naturalistic viewing is less established than that of theta. The robust feature across both bands is that the difference sits in slow frontocentral activity rather than over visual cortex.

That localization suggests the difference is not in raw stimulus processing, which would predict effects over visual cortex, but in higher-order attentional and integrative processes. Two observations support this. The evoked-magnitude control is an occipital, narrowband, primary visual measure, and it absorbs less of the frontocentral delta effect (7.7 to 11.6 percent) than of the whole-head broadband effect (8.6 to 18.1 percent) in every movie, which is what a genuinely non-sensory frontocentral effect would predict. The same ordering holds when quality-control covariates are added alongside it.

This inference concerns location rather than size. The frontocentral cluster is also where female recordings are most often interpolated, and in recordings with no cluster channel interpolated the effect is attenuated by 23 percent (Section 3.4.3), part of which reflects that stratum selecting cleaner recordings generally, so the asymmetry inflates the height of the effect at the cluster by an amount these data do not resolve precisely. It does not appear to determine the location: within the subsample for which per-channel interpolation records exist, restricting to recordings with no cluster channel interpolated leaves the maximum at E30 in both bands, though at that subsample size the maximum is unstable to removal of recordings by any rule, so this is consistent with rather than demonstrative of the location being independent of the asymmetry. What the data support is that the effect is frontocentrally weighted, is not visual-cortical, and peaks at a location the interpolation asymmetry does not appear to determine on the subsample where that can be tested. Taking the spectral, spatial and developmental characterizations together, the most parsimonious interpretation is that male participants maintain more uniform cross-subject patterns of slow frontocentral engagement during naturalistic viewing, most pronounced in pre-adolescence, with a minority share attributable to greater evoked response magnitude. We do not read “more uniform” as better or worse; both higher and lower ISC have been linked to different cognitive states in different contexts.

### 4.4 The CBCL Bound

That ISC predicted none of the four CBCL bifactor dimensions at controlled FDR, despite *n >* 1100 and robust age and sex signal in the same data, is a notable result. The bound is what makes it informative. Associations between parent-report psychometric instruments and neural measures are typically reported in the range *r* = 0.10 to 0.20; 13 of the 16 confidence intervals here do not reach the bottom of that range, and the three that do extend past it by 0.002 to 0.006. Nothing at |*β*| = 0.11 or above is compatible with these data at any of the 16 tests, and the bound is tighter still at the frontocentral cluster, where the widest interval reaches 0.086. The result is not that we failed to detect an association of the size the literature reports, but that we can exclude all but the very bottom edge of it.

Several considerations bear on interpretation. Grand-mean ISC compresses 129 channels and 117 to 203 seconds of dynamic response into a scalar, and finer features such as time-windowed ISC or subnetwork-specific ISC might carry signal that neither the grand mean nor the per-channel decomposition captures. CBCL bifactor scores are parent-reported and reflect general parental concern, which may not map onto neural measures during 2 to 3 minutes of viewing. Other measures from the same recordings, such as functional connectivity or source-localized power, might predict CBCL where ISC does not.

The result converges with two independent lines of evidence: null findings on HBN-EEG functional connectivity [13], and the 2025 NeurIPS EEG Foundation Challenge [17], in which over 1100 teams and participants predicted CBCL externalizing scores from HBN-EEG recordings across multiple paradigms and only three finished below the organizers’ threshold of 0.99, where 1.0 is the score obtained by predicting the mean; the winning normalized error was 0.978, a 2.2 percent reduction [18]. The organizers additionally report that samples in that challenge were not randomized, so contiguous trials from the same subject were exploitable, which makes 0.978 an optimistic figure. That a cohort-level summary statistic and short-snippet deep-learning embeddings converge on the same null suggests the difficulty is not specific to a feature representation or timescale but is a more general bound on what these metrics encode about parent-reported pediatric psychopathology. Future work will need feature representations capturing finer temporal, spatial or task-specific structure, and may need to consider whether the limitation lies in the EEG signal or in CBCL as a target.

### 4.5 Limitations

#### Reference and bad-channel detection

The common average reference was applied before bad-channel detection and was a plain average over all recorded channels rather than PyPREP’s robust reference, so channels later flagged as bad contributed to the average against which detection ran. This inflates flagging in noisy recordings, and recording noise differs by sex in this cohort, so part of the interpolation asymmetry may be introduced by the pipeline rather than present in the raw data. Two observations bound the consequence. The asymmetry persists among recordings in which the recovered reference channel was never flagged, so it is not an artifact of that channel alone; and the analyses in Section 3.4.3 condition on interpolation directly, so they bound its contribution to the sex effect whatever its origin. Re-running detection under a robust reference across the full cohort would settle the question and is the most direct methodological extension of this work.

#### Cohort composition and overlap

All findings derive from HBN. The direction of the sex effect has been reported in an independently recruited cohort [9], [10], which addresses independence, but neither that cohort nor ours is a population-representative typical-development sample, and replication in such a sample remains necessary. Our cohort also overlaps the replication sample of that report by an amount we cannot quantify; the release-exclusion analysis establishes that the effect does not depend on the overlapping subjects but does not identify them.

#### Sample composition confound

HBN is referred for clinical concern with a ~2:1 male skew enriched for ADHD-spectrum presentations. The low-symptom subset replication preserved the effect and attention CBCL is a covariate in the primary regression, but DSM diagnostic codes are absent from the public release, so residual confounding from undiagnosed clinical traits cannot be excluded.

#### Cross-sectional design

Age effects are inferred across age bins, not longitudinally. The inverted-U pattern is consistent with developmental change but may also reflect cohort effects.

#### Sex as recorded

Sex is a parent-reported binary field with no gender identity field, pubertal staging or hormonal measure available. The maturational interpretations in Section 4.1 concern developmental timing, which pubertal staging would index far better than chronological age and sex alone.

#### Clinical phenotype and target

CBCL scores are parent-reported and may not capture the aspects of phenotype most relevant to neural synchronization. The bifactor model has also been criticized on statistical grounds, since bifactor models fit correlated data well and the resulting factors may not correspond to underlying processes [31], [32]; the bound may therefore partially reflect target unreliability rather than predictor unreliability.

#### Stimulus set

Four clips of 2 to 3 minutes. Longer stimuli or systematic manipulation could identify which features drive the effect.

#### Evoked-magnitude control

Surround Suppression recordings were available for 354 of the 400 subjects in the band subsample. Availability was balanced by sex and unrelated to age, symptom scores or recording noise, but subjects without recordings had slightly lower grand-mean ISC (*p* = 0.015), so the subsample is marginally cleaner than the whole. The delta-cluster and broadband models rest on the same 353 subjects, so the two outcomes are not independent samples. SSVEP is a narrowband occipital measure of one evoked response, not a complete index of evoked response strength across the frequency range in which the ISC effect sits.

#### Location of the topographic peak

Per-channel interpolation records were retained only for the *n* = 150 subsample, so the test of whether the interpolation asymmetry places the t-statistic maximum at the cluster rests on 457 recordings rather than on the full cohort (Section 3.4.3). At that size the identity of the maximum is sensitive to which recordings are removed: under random removal at the same sex-specific rates, E30 is the maximum in 38 percent of delta draws and 64 percent of broadband draws. The clean-stratum outcome and the delta control outcome both fall within the range random removal produces, so the test supports only the weaker statement that removing the recordings the asymmetry affects does not move the maximum away from the cluster. It does not establish that the location is independent of recording quality in general. Retaining per-channel interpolation records for the full cohort would allow the test at full scale.

#### Analyses on subsamples

The band decomposition (*n* = 400), the ocular-removal and per-channel interpolation analyses (*n* = 150 each) and the stability comparison (*n* = 30) rest on subsamples, with correspondingly wider intervals. The same-sex template analysis is reported at both scales: Test A on the balanced *n* = 400 subsample and Test B on the full cohort. The ocular re-fit resampled to 200 Hz before independent component analysis rather than after, a tractability tradeoff; it reproduced the published cleaning to a median per-channel correlation above 0.97 and recovered the effect under test.

### 4.6 Conclusion

We characterized EEG inter-subject correlation in the largest pediatric naturalistic-stimulus cohort to date and confirmed a low-frequency frontocentral male advantage, largest in delta, peaking in pre-adolescence, in the same direction as a previously reported effect that was marginal at the sample sizes then available. The effect survived three demographic and clinical robustness analyses, three analyses targeting the ISC computation and the measurement, and exclusion of the data releases overlapping that earlier report. Differential channel interpolation attenuates the effect, by approximately 10 percent under covariate adjustment and by 23 percent in recordings with no cluster channel interpolated, but does not explain it. Evoked response magnitude, previously proposed as an alternative account, is confirmed to differ by sex but carries 8.6 to 18.1 percent of the effect. We additionally find that ISC does not predict CBCL bifactor scores at *n >* 1100 at any of three feature resolutions, with confidence intervals excluding associations larger than |*β*| = 0.106. Replication in independently recruited typical-development samples remains necessary to discriminate among causal interpretations, but pediatric EEG ISC during naturalistic viewing is sex-dependent in a non-trivial way during the developmental window of greatest interest.

## Author Contributions (CRediT)

E. Tomar: Conceptualization, Methodology, Software, Formal analysis, Investigation, Data curation, Visualization, Writing (original draft), Writing (review and editing), Project administration. A. Silvan: Resources, Writing (review and editing). E. Y. Fu: Supervision, Validation (results and analysis), Writing (review and editing). Y. Xiong: Writing (review and editing). C. W. Do: Supervision.

## Acknowledgment

Portions of the analysis code were developed, and portions of the manuscript drafted and edited, with the assistance of large language model tools (Anthropic Claude). All analyses, results, and claims were verified by the authors, who take full responsibility for the content. The authors thank the Child Mind Institute and the Healthy Brain Network participants and their families for the public availability of the HBN-EEG dataset.

