## Supplementary Materials for "Sex differences in pediatric EEG inter-subject correlation during naturalistic movie watching: a large-scale characterization of the Healthy Brain Network EEG dataset"

### S1 Preprocessing Pipeline Validation

The `v3concat` pipeline was developed iteratively, with two earlier iterations (v1, v2) tested on a subset of subjects before convergence. Validation of the final pipeline was performed via a single-subject pilot in which `v3concat` output was compared to per-movie ICA fits on the same data. The resulting cleaned signals showed median per-channel correlation of 0.815 to 0.869 across the four movies, with a long lower tail (per-channel minima of $-0.71$ to 0.12). Agreement between the two decompositions is therefore broad at the typical channel but not uniform across channels; the joint fit additionally provides a single uniform exclusion list per subject.

This pilot rests on one subject and should be read as a sanity check on the implementation rather than as a characterization of how the two approaches differ. The quantitative comparison that carries the methodological argument is the decomposition stability analysis in Section S6, which uses 30 subjects and 20 initializations per condition. Extending the signal-level agreement analysis, and the downstream consequences of the cleaning choice, across a larger cohort is reserved for future methods-focused work.

### S2 Supplementary Figures

The following figures accompany the robustness and characterization analyses in the main text and are generated by `scripts/figures/supplementary.py`, except Figure S2, which is generated by `scripts/figures/fig_s2_qc_tiers.py`.

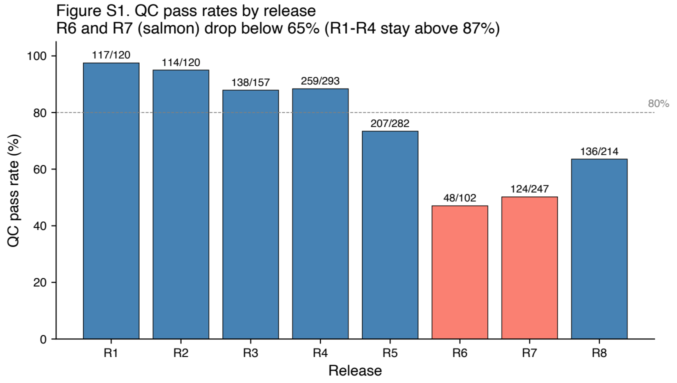


Fig. S1. Quality-control pass rates by HBN release, showing the lower pass rates in R6 and R7 noted in Section 2.5 of the main text.

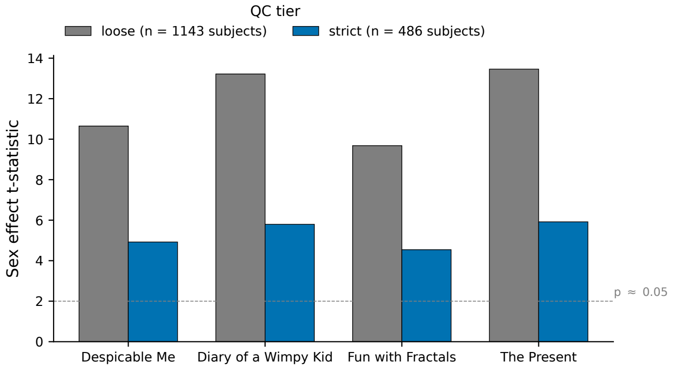


Fig. S2. Sex effect t-statistic at two QC tiers, the full cohort (1143 subjects) and the strict QC subset (486 subjects). Bars are two-sample t-statistics, which decline partly through power loss at smaller $n$; the underlying effect size (Cohen's $d$) shrinks by a smaller amount (about 27%), whose source is analyzed in Section 3.4.2 of the main text. The effect direction is preserved throughout.

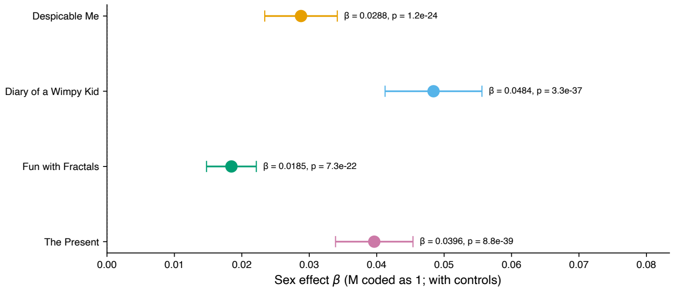


Fig. S3. Forest plot of the marginal per-movie sex coefficients from the fully adjusted model, accompanying the sex-by-movie interaction test in Section 3.4.8 of the main text. The direction is uniform across movies while the magnitude varies.

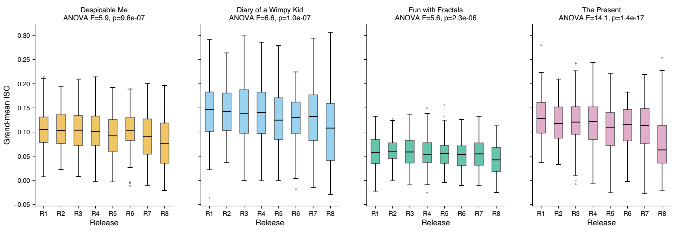


Fig. S4. Distribution of grand-mean ISC by release for each movie, accompanying the release-effect control in the statistical analyses.

### S3 Validity Analyses

This section reports the full numerical results of the methodological validity analyses summarized in Section 3.4.3 of the main text: robustness of the male-greater-than-female ISC effect to same-sex and size-matched leave-one-out templates, controls for ocular artifact, and the channel interpolation analysis. The fourth validity analysis, on evoked response magnitude, is reported in Section S4.

### S3.1 Same-Sex and Size-Matched Templates

The leave-one-out template is the mean of all other subjects, so the 2:1 male majority makes it male-weighted. Two tests address this. Test A recomputes the theta frontocentral effect on the balanced $n = 400$ subsample, where the two same-sex templates are already size-matched at about 199 subjects each. Test B operates on the full cohort of 1143, where the male template pools more subjects and is therefore less noisy; the male template is subsampled to the female sample size in each movie (378, and 377 for The Present) and averaged over 50 random draws (seed 42) so both templates pool an equal number of subjects. Test A was run in theta, the band in which this gate was originally specified; Test B is broadband and therefore dominated by the low-frequency bands that carry most of the ISC signal. Table S1 reports both. The effect strengthens rather than weakens in every case. In Test A the same-sex template lowers female theta cluster ISC from 0.01502 to 0.00898 while leaving male ISC almost unchanged (0.03126 to 0.03030). Some asymmetry is expected from template noise alone, because the group with lower baseline ISC is more noise-limited and loses proportionally more when the template shrinks; the observed 40 percent against 3 percent is substantially larger than that mechanism predicts.

### S3.2 Ocular Artifact Controls

Because the HBN public release contains no eye-tracking, these controls are entirely EEG-internal. The ocular signal is the set of independent components that ICALabel classifies as eye blink. Two tiers are reported: spatial and spectral characterization that needs no re-fitting (Tables S2 and S3), and a direct pre versus post ocular-removal test that re-fits the ICA per subject (Table 5 of the main text and Table S4 below).

#### *S3.2.1 Spatial and Spectral Tests*

An ocular source peaks at the electrodes nearest the eyes and falls off with distance; a frontal-midline neural source peaks frontocentrally. Table S2 shows the theta sex-effect t-statistic peaks on the frontocentral cluster and is null at the periocular electrodes, the reverse of an ocular gradient, in every movie. Electrode groups were derived from the GSN-HydroCel-129 montage geometry: the periocular ring is the ten electrodes nearest the two montage-derived orbit points (E126, E127, E25, E8, E17, E21, E14, E1, E32, E125), and the frontopolar-midline set is the anterior near-midline electrodes (E10, E11, E14, E15, E16, E17, E18, E21). Three electrodes (E14, E17, E21) fall in both groups; both serve as non-frontocentral comparison regions, so the overlap does not affect the contrast against the frontocentral cluster.

Blink residual is spectrally concentrated in delta (1 to 4 Hz). Table S3 reports the sex effect by band. It is larger in delta than in theta, the low-frequency profile blink residual would also show, but it peaks on the same frontocentral cluster (delta mean $t = 9.23$ vs periocular mean $t = 1.55$) rather than periocularly, a low-frequency neural topography rather than an ocular one. The full frontocentral cluster profile in Hedges $g$ is delta 1.348, theta 0.889, alpha 0.196, beta 0.133, so the delta effect exceeds the theta effect by a factor of 1.52 at the cluster, and the higher bands that would carry broadband saccadic-spike artifact (alpha, beta) are essentially null.

#### *S3.2.2 Pre Versus Post Ocular-Component Removal*

The ICA was re-fit per subject on a balanced subsample ($n = 150$, 75 male, 75 female; 149 for The Present, which excludes one recording below the 200 s minimum duration), and ISC was recomputed under three conditions produced from one fit by varying the exclusion list: A removes all artifact components including ocular (the published condition), B retains the ocular components and removes the rest, and C removes nothing. For tractability the re-fit resampled to 200 Hz before ICA; condition A regenerated this way reproduced the published cleaning to a median per-channel correlation of 0.978 to 1.000 across five check subjects. Table 5 of the main text shows the frontocentral effect is invariant across all three conditions in both bands.

The same manipulation moves ISC strongly at the periocular electrodes but not at the frontocentral cluster, a double dissociation confirming the ocular components reach the eyes but not the cluster. In the subset of subjects with at least one eye blink component ($n = 112$; 62 male, 50 female), the isolated ocular signal carries no detectable frontocentral sex difference (delta Hedges $g = -0.056$, $p = 0.77$; theta $g = 0.009$, $p = 0.96$), and the sex effect is undiminished after controlling for it. The dissociation holds in each of the four movies individually, ranging from 9.9 to 49.7 fold within movies against 16 fold pooled. As a further check on movement and drift contamination, per-recording amplitude (median per-channel standard deviation) correlates negatively with delta ISC ($r = -0.26$, $p < 10^{-6}$), the only band showing a significant correlation, which is the opposite of what movement or drift contamination would produce. Table S4 reports the dissociation and the covariate regression.

### S3.3 Channel Interpolation

Channels flagged as bad by PyPREP were spherically interpolated from their neighbors, so a sex difference in interpolation frequency could bias ISC. Table S5 reports interpolated channel counts by sex and their association with grand-mean ISC. Counts were retained for the full cohort; per-channel interpolation identities were retained only for the balanced $n = 150$ subsample, so the cluster-level analysis in Table S6 is restricted to that subsample.

The association between interpolation count and ISC is negative in every movie, so interpolation count behaves as a proxy for recording noise rather than as a source of smoothing-induced inflation. Because female recordings carry more interpolated channels, the asymmetry depresses female ISC and widens the observed sex gap.

In the $n = 150$ subsample, interpolation at the frontocentral cluster lowers cluster delta ISC from 0.080 to 0.042, a reduction of 47.8 percent. Restricting to recordings in which

TABLE S1

Robustness to same-sex and size-matched templates. Welch $t$ and small-sample Hedges $g$, males vs females. Test A: theta frontocentral cluster (E30, E36, E37) on the balanced $n = 400$ subsample. Test B: broadband (1 to 20 Hz) on the full cohort ($n = 1143$), male template size-matched to the per-movie female sample size (377 or 378) over 50 draws.

| Analysis | Template | Δ (M−F) | Welch $t$ | Hedges $g$ |
|---|---|---|---|---|
| *Test A: theta frontocentral cluster ($n = 400$)* | | | | |
| | pooled (mixed-sex) | +0.01624 | 8.90 | 0.889 |
| | same-sex | +0.02132 | 12.77 | 1.274 |
| *Test B: broadband whole-head grand mean ($n = 1143$)* | | | | |
| | pooled (2:1 skew) | +0.03408 | 13.50 | 0.873 |
| | same-sex matched | +0.03966 | 18.44 | 1.140 |
| *Test B: broadband frontocentral cluster ($n = 1143$)* | | | | |
| | pooled (2:1 skew) | +0.06011 | 21.87 | 1.376 |
| | same-sex matched | +0.06742 | 31.84 | 1.751 |

TABLE S2

Theta sex-effect t-statistic (males minus females) by electrode group, per movie. The effect peaks frontocentrally and is null at the periocular electrodes nearest the eyes.

| Electrode group | DM | DoaWK | FwF | TP | Mean |
|---|---|---|---|---|---|
| Frontocentral (E30, E36, E37) | 5.53 | 5.87 | 3.97 | 5.66 | 5.26 |
| Periocular ring (10) | 0.76 | 0.59 | 0.37 | 0.56 | 0.57 |
| Frontopolar midline (8) | 1.74 | 1.07 | −0.31 | 0.50 | 0.75 |

TABLE S3

Sex effect on band-specific ISC, balanced $n = 400$ subsample. Welch $t$ and Hedges $g$, males vs females, for the frontocentral cluster and the whole-head grand mean.

| Band | Region | mean M | mean F | Δ | $t$ | $g$ |
|---|---|---|---|---|---|---|
| delta | frontocentral | 0.10735 | 0.04863 | +0.05871 | 13.51 | 1.348 |
| theta | frontocentral | 0.03126 | 0.01502 | +0.01624 | 8.90 | 0.889 |
| delta | whole-head | 0.10839 | 0.07430 | +0.03409 | 9.02 | 0.900 |
| theta | whole-head | 0.02934 | 0.02226 | +0.00708 | 4.65 | 0.464 |

TABLE S4

Top: change in pooled delta ISC when ocular components are retained (B) versus removed (A), by region. Bottom: regression of condition-A frontocentral delta ISC on sex (male coded 1), with and without the isolated ocular ISC as a covariate.

| *Double dissociation (delta, pooled ISC)* | | | |
|---|---|---|---|
| Region | ISC (A) | ISC (B) | change A to B |
| Frontocentral cluster | 0.07123 | 0.07232 | +1.5% |
| Periocular ring | 0.06457 | 0.08032 | +24.4% |

| *Covariate regression (A frontocentral delta ISC)* | | | | |
|---|---|---|---|---|
| Model | sex $\beta$ | SE | $t$ | $p$ |
| sex only | +0.03588 | 0.00716 | 5.01 | 2.06e−06 |
| sex + ocular-only | +0.03629 | 0.00706 | 5.14 | 1.23e−06 |

no cluster channel was interpolated ($n = 266$ male, 191 female), the frontocentral sex effect is delta Hedges $g = 0.585$ ($p = 8.5 \times 10^{-10}$) and theta $g = 0.341$ ($p = 3.6 \times 10^{-4}$), against delta $g = 0.975$ in the interpolated stratum and $g = 0.760$ pooled across both, an attenuation of 23 percent. All three are recording-level figures; the value of 1.133 in Table 5 of the main text is computed at the subject level and is not their comparator. A cluster bootstrap (2000 resamples, seed 42) resampling subjects with replacement, with the ratio computed at the recording level so that within-subject clustering is respected, places the clean-to-pooled ratio at 0.770 with a 95 percent interval of 0.582 to 0.911. These stratum effect sizes are computed on the per-subject re-fit used for the ocular conditions, not on the primary preprocessing

TABLE S5
Interpolated channels per recording by sex, full cohort, and the association between interpolation count and grand-mean ISC. Counts are out of 129 channels. Welch $t$ is males versus females; a negative $t$ indicates more interpolation in female recordings.

| Scope | M mean (SD) | F mean (SD) | Welch $t$ | $p$ | $r$ with ISC |
|---|---|---|---|---|---|
| Pooled | 21.86 (8.91) | 23.91 (9.50) | −7.00 | < 0.001 | −0.328 |
| Despicable Me | 21.67 (8.95) | 23.15 (9.26) | −2.56 | 0.011 | −0.356 |
| Diary of a Wimpy Kid | 21.31 (8.68) | 24.04 (9.15) | −4.83 | < 0.001 | −0.343 |
| Fun with Fractals | 23.93 (9.21) | 25.51 (9.98) | −2.58 | 0.010 | −0.294 |
| The Present | 20.52 (8.42) | 22.93 (9.41) | −4.22 | < 0.001 | −0.317 |

All ISC correlations significant at $p < 1 \times 10^{-23}$.

TABLE S6
Top: frequency of frontocentral cluster channel interpolation by sex, balanced $n = 150$ subsample (599 recordings). Bottom: primary per-movie regression with and without interpolation count as an additional covariate, full cohort. Coefficients without the covariate reproduce the published values in Table 4 of the main text exactly.

| *Cluster interpolation frequency ($n = 150$)* | | | | | |
|---|---|---|---|---|---|
| Electrode | Male | Female | Male $n$ | Female $n$ | Fisher $p$ |
| E30 | 5.00% | 12.04% | 15/300 | 36/299 | 0.002 |
| E36 | 5.33% | 17.06% | 16/300 | 51/299 | < 0.001 |
| E37 | 5.33% | 20.07% | 16/300 | 60/299 | < 0.001 |
| Any of the three | 11.33% | 36.12% | 34/300 | 108/299 | < 0.001 |
| *Sex coefficient with and without interpolation covariate ($n = 1143$)* | | | | | |
| Movie | $\beta$ without | $\beta$ with | $p$ with | $\eta_p^2$ with | Retained |
| Despicable Me | +0.0288 | +0.0261 | 3.0e−23 | 0.085 | 90.8% |
| Diary of a Wimpy Kid | +0.0484 | +0.0425 | 1.5e−31 | 0.115 | 87.7% |
| Fun with Fractals | +0.0185 | +0.0167 | 1.1e−19 | 0.071 | 90.3% |
| The Present | +0.0396 | +0.0359 | 2.0e−34 | 0.126 | 90.6% |

pipeline that yields Table S7 below. On the same recordings and the same stratification the primary pipeline gives clean $g = 0.638$, interpolated 1.167 and pooled 0.838, with the same ordering and a clean-to-pooled ratio of 0.762 against 0.770, so the two bases agree on the structure and differ on the level.

Two removal controls separate the effect of cluster interpolation from that of general recording quality. Removing recordings at the same sex-specific rates but at random gives mean $g = 0.756$ over 200 draws (95 percent interval 0.677 to 0.852), reproducing none of the attenuation, so the attenuation is not a consequence of asymmetric sample loss alone. Removing them at the same rates but ranked on interpolation outside the cluster gives $g = 0.655$, between the pooled and clean values, with a bootstrap interval of 0.411 to 0.916 spanning both, so the share attributable to cluster interpolation specifically is not resolvable at this subsample size. The same limit applies to the region comparison: the clean-to-pooled ratio is 0.770 at the cluster (95 percent CI 0.582 to 0.911) against 0.599 for the whole-head delta effect (0.203 to 0.865), with the interval on their difference including zero (−0.486 to +0.030). Both ratios exclude zero, so a male advantage survives in the clean stratum in both regions.

The whole-head interpolation asymmetry in this subsample is 24.39 against 20.37 channels, a ratio of 1.2, against 3.2 at the cluster.

Because the common average reference precedes bad-channel detection, the recovered Cz channel is itself screened, and it is flagged far more often in female than in male recordings (35.45 against 6.33 percent in the n = 150 subsample, Fisher $p = 1.3 \times 10^{-19}$), consistent with its post-reference trace being the negative mean of all other channels and therefore a readout of montage-wide noise. The sex difference in flagged channels is not created by this mechanism: among recordings in which Cz was never flagged it remains 22 against 19 channels (Mann-Whitney $p = 3.9 \times 10^{-3}$). Excluding Cz from the 129-channel grand mean changes the four primary sex coefficients by 0.26 to 0.69 percent, in every case a reduction.

Two limits apply: the cluster percentages rest on 150 subjects rather than the full cohort, and the regression covariate is a whole-head count used as a proxy for cluster-specific interpolation.

### S4 Evoked Response Magnitude

This section reports the full results of the evoked-magnitude control summarized in Section 3.4.4 of the main text, which reproduces the steady-state visual evoked potential (SSVEP) control of Petroni et al. (2018) on the HBN Surround Suppression task. References are as numbered in the main text.

TABLE S7
Per-channel sex-effect topography under stratification on cluster interpolation. Recording level, pooled across movies, computed on the primary preprocessing pipeline. Map R is all 599 recordings of the balanced $n = 150$ subsample. Map C restricts to recordings in which no frontocentral cluster channel was interpolated. Map Y removes the same numbers of recordings as map C but ranked on interpolation outside the cluster. FULL is the unstratified reference: the $n = 400$ band subsample for delta and the full cohort for broadband. Cluster columns give the rank of that channel in the map with its t-statistic. Spatial $r$ is the correlation across all 129 channels against map R of the same band.

| Band | Map | $n_M$ | $n_F$ | max-$t$ ch | max $t$ | E30 | E36 | E37 | spatial $r$ |
|---|---|---|---|---|---|---|---|---|---|
| delta | R | 300 | 299 | E30 | 9.61 | 1 (9.61) | 3 (9.36) | 4 (9.17) | 1.000 |
| delta | C | 266 | 191 | E30 | 6.76 | 1 (6.76) | 6 (6.00) | 8 (5.67) | 0.980 |
| delta | Y | 266 | 191 | E29 | 7.44 | 3 (6.88) | 2 (7.35) | 5 (5.96) | 0.982 |
| delta | FULL | 799 | 799 | E30 | 17.43 | 1 (17.43) | 3 (16.70) | 2 (16.72) | 0.964 |
| broadband | R | 300 | 299 | E30 | 10.96 | 1 (10.96) | 3 (10.58) | 2 (10.65) | 1.000 |
| broadband | C | 266 | 191 | E30 | 7.89 | 1 (7.89) | 5 (6.90) | 2 (7.27) | 0.984 |
| broadband | Y | 266 | 191 | E36 | 8.21 | 2 (8.20) | 1 (8.21) | 5 (7.60) | 0.982 |
| broadband | FULL | 3057 | 1511 | E30 | 31.02 | 1 (31.02) | 2 (29.10) | 4 (28.48) | 0.951 |

Random-removal control (map X), 200 draws removing recordings at the same sex-specific rates as map C: E30 is the maximum-t channel in 77 of 200 delta draws, where the modal channel is E29 at 84 of 200, and in 127 of 200 broadband draws.

#### S4.1 Measure Validation

Three checks establish that the extracted quantity is a stimulus-driven SSVEP rather than noise at the flicker frequency. First, the group-mean signal-to-noise ratio at 25 Hz peaked at electrode E76, with all five analysis electrodes (E76, E75, E71, E72, E77) posterior, as required for a primary visual response. Second, SNR scaled with foreground contrast, sitting at the noise floor in the no-flicker condition (Table S8, upper block); the paired within-subject excess of high-contrast (at or above 60 percent) over zero-contrast SNR was $+0.2743$ (SD 0.3569, $t(353) = 14.46$, Cohen's $d = 0.77$), positive in 85.0 percent of subjects. Third, split-half reliability across odd and even trials was $r = 0.891$, or $r = 0.943$ after Spearman-Brown correction, so the per-subject estimate is reliable enough to serve as a covariate.

Trial rejection used a robust global criterion computed across channels rather than an any-channel threshold. With 129 channels and no prior artifact removal, an any-channel rule is triggered by any persistently bad electrode on essentially every trial; simulation indicates that five bad channels are sufficient to reject all trials under such a rule. Because bad-channel counts differ by sex in this cohort (Table S5), that failure mode would have biased the covariate on the variable under test. Under the robust criterion, median trial retention was 87.5 percent and no subject was excluded.

#### S4.2 Sample and Selection

Of the balanced $n = 400$ subsample, 354 have Surround Suppression recordings in the public release, and extraction excluded no further subjects. The 46 subjects without recordings did not differ from the 354 with them on sex (178 male and 176 female with, 22 male and 24 female without; Fisher exact $p = 0.876$, odds ratio 1.10), age ($p = 0.33$), general psychopathology factor ($p = 0.68$), attention ($p = 0.39$), or recording noise ($p = 0.73$). They did have slightly lower grand-mean ISC (0.0774 against 0.0940, Welch $t = 2.49$, $p = 0.015$), so the subsample used here is marginally cleaner than the band subsample as a whole.

#### S4.3 Robustness Across SSVEP Measures

Table 6 of the main text reports the effect of control using the incoherent measure, which matches the per-trial averaging used in the original report. Substituting either coherently averaged measure gives the same conclusion. On broadband grand-mean ISC, the sex coefficient retained 81.8, 92.3, 86.9 and 89.9 percent under the coherent all-trials measure, and 78.1, 92.1, 84.3 and 87.4 percent under the coherent high-contrast measure, for Despicable Me, Diary of a Wimpy Kid, Fun with Fractals and The Present respectively. All coefficients remained significant at $p < 5 \times 10^{-6}$ in every movie under every measure.

### S5 Release Exclusion Sensitivity

The replication cohort of Petroni et al. ($n = 303$) was drawn from HBN, so the present cohort overlaps theirs by an amount we cannot determine, since subject identifiers for their sample are not published. Because their sample predates the later HBN releases, we re-ran the primary regression with the earliest releases excluded. Table S9 reports all four exclusion levels. The sex effect persists in every movie at every level, at $p < 1 \times 10^{-20}$ throughout, with a third of the cohort removed at the strictest level. Level (a) reproduces Table 4 of the main text and serves as a check on the procedure.

Coefficients do not decrease under exclusion. We do not interpret the direction of that change, because release composition differs on several dimensions and release is already modeled as a fixed effect in the primary regression; the analysis establishes only that the effect does not depend on the overlapping subjects.

### S6 Decomposition Stability

Table S10 reports the full per-condition results of the ICA stability comparison summarized in the main text. $I_q$ is the ICASSO cluster quality index, computed over 20 fits per condition per subject on 30 subjects, with the same 20 random initializations used in every condition so that comparisons are paired. Fits were performed at the native

TABLE S8

Evoked response magnitude: measure validation and sex difference. Upper block: group-mean 25 Hz signal-to-noise ratio by foreground contrast, over the five analysis electrodes, complete-case subjects only. Middle block: sex difference in each of the three SSVEP measures. Lower block: correlation between the primary (incoherent) SSVEP measure and grand-mean ISC, per movie. Values for the corresponding analyses in Petroni et al. are given for comparison where available.

| *Contrast response function ($n = 354$)* | | | | | | |
|---|---|---|---|---|---|---|
| Foreground contrast | SNR | | excess over 0% | | | |
| 0% (no flicker) | 1.0008 | | n/a | | | |
| 30% | 1.1620 | | +0.1612 | | | |
| 60% | 1.2444 | | +0.2436 | | | |
| 100% | 1.3056 | | +0.3049 | | | |
| *Sex difference in SSVEP magnitude* | | | | | | |
| Measure | mean M | mean F | SD M | SD F | Welch $t$ | Cohen $d$ |
| Incoherent (primary) | 1.2377 | 1.1174 | 0.2552 | 0.1721 | 5.21 | 0.55 |
| Coherent, all trials | 2.4525 | 1.6023 | 1.7090 | 1.0451 | 5.65 | 0.60 |
| Coherent, contrast $\geq$ 60% | 2.6500 | 1.7380 | 1.9275 | 1.1574 | 5.40 | 0.57 |
| *SSVEP to ISC correlation (incoherent measure)* | | | | | | |
| Movie | $r$ | | $p$ | | $n$ | |
| Despicable Me | +0.245 | | 3.2e−06 | | 354 | |
| Diary of a Wimpy Kid | +0.202 | | 1.3e−04 | | 354 | |
| Fun with Fractals | +0.247 | | 2.5e−06 | | 354 | |
| The Present | +0.251 | | 1.9e−06 | | 353 | |

Petroni et al. reported no significant sex difference in SSVEP magnitude ($F(1, 106) = 3.3$, $p = 0.08$) and an ISC to SSVEP correlation of $r = 0.41$ ($p = 0.0001$, $N = 84$). All sex differences above are significant at $p < 4 \times 10^{-7}$.

TABLE S9

Sex effect on grand-mean ISC under progressive exclusion of the earliest HBN releases, which are those most likely to overlap the replication cohort of Petroni et al. Level (a) includes all releases and reproduces Table 4 of the main text. $n$ is the per-movie regression sample; subject counts are the number of unique subjects retained. All coefficients positive (males greater than females) at every level.

| Exclusion level | Movie | $n$ | $\beta$ | $t$ | $p$ | $\eta_p^2$ |
|---|---|---|---|---|---|---|
| *(a) all releases, 1143 subjects* | | | | | | |
| | Despicable Me | 1132 | +0.02878 | 10.50 | 1.2e−24 | 0.0898 |
| | Diary of a Wimpy Kid | 1132 | +0.04843 | 13.23 | 3.3e−37 | 0.1355 |
| | Fun with Fractals | 1132 | +0.01846 | 9.81 | 7.3e−22 | 0.0794 |
| | The Present | 1128 | +0.03964 | 13.54 | 8.8e−39 | 0.1416 |
| *(b) excluding R1, 1026 subjects* | | | | | | |
| | Despicable Me | 1017 | +0.02950 | 10.11 | 6.1e−23 | 0.0926 |
| | Diary of a Wimpy Kid | 1017 | +0.05032 | 13.01 | 6.9e−36 | 0.1446 |
| | Fun with Fractals | 1017 | +0.01887 | 9.57 | 7.9e−21 | 0.0838 |
| | The Present | 1013 | +0.04048 | 13.02 | 6.4e−36 | 0.1453 |
| *(c) excluding R1 and R2, 912 subjects* | | | | | | |
| | Despicable Me | 907 | +0.03161 | 10.08 | 1.1e−22 | 0.1022 |
| | Diary of a Wimpy Kid | 907 | +0.05464 | 13.28 | 7.7e−37 | 0.1648 |
| | Fun with Fractals | 907 | +0.02040 | 9.70 | 3.1e−21 | 0.0954 |
| | The Present | 903 | +0.04268 | 12.82 | 1.3e−34 | 0.1559 |
| *(d) excluding R1 to R3, 774 subjects* | | | | | | |
| | Despicable Me | 771 | +0.03343 | 9.79 | 2.2e−21 | 0.1122 |
| | Diary of a Wimpy Kid | 771 | +0.05612 | 12.64 | 2.2e−33 | 0.1741 |
| | Fun with Fractals | 771 | +0.02167 | 9.68 | 5.8e−21 | 0.1100 |
| | The Present | 767 | +0.04437 | 12.29 | 9.4e−32 | 0.1668 |

500 Hz sampling rate. The two duration-matched conditions both use 85000 samples: concat170 is a contiguous 170 s window drawn from the concatenated stream and therefore spans a clip boundary, while clip170 is a single 170 s clip.

The two duration-matched conditions answer what the concatenation contributes. At 85000 samples each, concat170 reaches mean $I_q$ 0.730 against 0.698 for clip170, so at matched sample volume the stream spanning a clip boundary is the more reproducible of the two and the benefit is attributable to content heterogeneity rather than to volume alone. Sample volume is not irrelevant: the full concatenation at 325351 samples exceeds concat170 at 85000. What the comparison rules out is that volume is the whole explanation.

The rightmost column reports the proportion of fits that reached the 500-iteration limit rather than converging. Non-convergence is near universal in every condition, including the full concatenation, so the benefit of concatenation is to the reproducibility of the recovered decomposition and not to convergence.

TABLE S10
ICA decomposition stability by condition, 30 subjects, 20 initializations per condition. $I_q$ is the ICASSO cluster quality index; higher is more reproducible across initializations. Components with $I_q \geq 0.8$ are counted out of 20. Brain $I_q$ restricts to components classified as brain by ICALabel.

| Condition | samples | mean $I_q$ | SD | $I_q \geq 0.8$ (of 20) | brain $I_q$ | reached max iter |
|---|---|---|---|---|---|---|
| concatenated | 325351 | 0.759 | 0.102 | 8.6 | 0.739 | 98.7% |
| clip, Fun with Fractals | 81496 | 0.723 | 0.109 | 6.6 | 0.700 | 99.8% |
| clip, The Present | 99882 | 0.696 | 0.096 | 4.5 | 0.687 | 92.7% |
| clip, Despicable Me | 85275 | 0.695 | 0.108 | 4.7 | 0.684 | 99.5% |
| clip, Diary of a Wimpy Kid | 58699 | 0.691 | 0.107 | 5.4 | 0.685 | 98.0% |
| concat170 (duration-matched) | 85000 | 0.730 | 0.101 | 6.3 | 0.714 | 99.2% |
| clip170 (duration-matched) | 85000 | 0.698 | 0.104 | 5.2 | 0.673 | 99.7% |
